# How Modulation of the Tumor Microenvironment Drives Cancer Immune Escape Dynamics

**DOI:** 10.1101/2024.08.16.608314

**Authors:** Pujan Shrestha, Zahra S. Ghoreyshi, Jason T. George

## Abstract

Metastatic disease is the leading cause of cancer-related death, despite recent advances in therapeutic interventions. Prior modeling approaches have accounted for the adaptive immune system’s role in combating tumors, which has led to the development of stochastic models that explain cancer immunoediting and tumor-immune co-evolution. However, cancer immunemediated dormancy, wherein the adaptive immune system maintains a micrometastatic population by keeping its growth in check, remains poorly understood. Immune-mediated dormancy can significantly delay the emergence (and therefore detection) of metastasis. An improved quantitative understanding of this process will thereby improve our ability to identify and treat cancer during the micrometastatic period. Here, we introduce a generalized stochastic model that incorporates the dynamic effects of immunomodulation within the tumor microenvironment on T cell-mediated cancer killing. This broad class of nonlinear birth-death model can account for a variety of cytotoxic T cell immunosuppressive effects, including regulatory T cells, cancer-associated fibroblasts, and myeloid-derived suppressor cells. We develop analytic expressions for the likelihood and mean time of immune escape. We also develop a method for identifying a corresponding diffusion approximation applicable to estimating population dynamics across a wide range of nonlinear birth-death processes. Lastly, we apply our model to estimate the nature and extent of immunomodulation that best explains the timing of disease recurrence in bladder and breast cancer patients. Our findings quantify the effects that stochastic tumor-immune interaction dynamics can play in the timing and likelihood of disease progression. Our analytical approximations provide a method of studying population escape in other ecological contexts involving nonlinear transition rates.

---

Metastatic disease remains the top contributor to cancer mortality despite the continued development of improved therapies. A large body of stochastic modeling and empirical work have established mechanisms of primary cancer progression via pre-existent and acquired resistance, through which therapeutic intervention commonly results in the evolutionary selection of resistant populations (1–5). Recent strides in cancer treatment have been made by enhancing the anti-tumor capacity of the adaptive immune system (6). In this context, experimental investigation and complementary stochastic models have been developed to study cancer immunoediting, explain cancer incidence (7–10), and correlate observed evolutionary patterns of disease with tumor-immune co-evolution (11–17).

Immune-mediated cancer dormancy, in contrast, represents a distinct and understudied problem related to cancer metastatic control. Early on, tumor cells extravasate from the primary disease site into circulation, where they can then ultimately settle at distant metastatic locations (18). The maintenance of actively growing cell populations at small sizes with balanced cell death represents one mechanism through which time to metastasis can be substantially prolonged (19, 20).

As with primary disease, micrometastatic cancer populations are also subjected to immunoediting by the adaptive immune system, wherein appreciable disease proceeds through elimination, equilibrium, and escape phases (21). Disruptions to balanced cancer growth and T cell killing in immunemediated dormancy can result in clinically appreciable disease. In two unfortunate clinical cases, kidney transplant recipients from a shared donor with prior history of primary melanoma developed either secondary melanoma at the kidney site or subcutaneous melanoma, likely as a result of their transplant-specific immunosuppressive regimen (22). In separate studies, carcinogen-injected mice that did not develop progressive malignancy with stable masses at the injection site were treated with control antibody or antibody-depleting T cells (23). In contrast with the control group, mice treated with T cell immune-depleting therapy developed progressive malignancy, and a repeated experiment with mice lacking adaptive immunity resulted in very few late-forming tumors, further implicating the role of T cell immunity in the longterm maintenance of cancer equilibrium (24).

Anti-tumor T cell responses can also be significantly affected by the presence of immunomodulatory elements in the tumor microenvironment. Immunomodulation is complex and involves a variety of chemical and cellular inputs. For example, cancer and stromal cells deprive T cells of nutrients, thereby encouraging exhaustion (25). Regulatory T cells (Tregs) and cancer associated fibroblasts (CAFs) accumulate in the microenvironment and inhibit T cells directly and indirectly (26–30). MDSCs expand and contract with tumor size, and have been shown to sequester cysteine from T cells and contribute to the presence of Tregs (31–35).

The near-impossibility of long-term clinical studies tracking micrometastatic initiation and growth necessitates a mathematical modeling framework capable of evaluating the timing and likelihood of cancer immune-mediated dormant resurgence as a function of dynamical signals and cell types in the metastatic immune microenvironment. Prior models have been proposed to study immune-mediated dormancy, including ODE models of tumor-immune interactions (36, 37). These models, while capable of describing mean behavior, characterize cell sizes on a continuum, and so scenarios involving equilibrium population sizes that are close to zero neglect the nontrivial extinction probability of this absorbing state and thus susceptibility to stochastic fluctuations. Their inability to resolve the distributional behavior of zero net-growth rates with strictly positive cancer cell division make them ill-equipped to differentiate balanced birth and death from zero birth and death, particularly in the regime of small population size.

To address the above questions and current limitations, we develop a generalized stochastic modeling framework that incorporates the effects of dynamical immunomodulatory cells present in the microenvironment. We construct a nonlinear birth-death model, wherein immunomodulatory regulation of effector T cells impairs their cancer killing ability. When applied to clinical data on cancer progression, our model predicts that the timing and likelihood of breast and bladder cancer is best explained by a microenvironment that varies dynamically with respect to the tumor burden. We also develop a diffusion approximation sufficiently general to handle reasonable assumptions on nonlinear birth-death processes, which we anticipate will be of broad use for additional applications that involve the description of nontrivial per-capita death rates.

## Material and Methods

### Model Development

Our model considers a small dividing cancer population experiencing linear growth and active killing by cytotoxic (CD8+) T cells . We assume that the cancer cells exhibit exponential growth and that robust immune responses are capable of imparting a high per-cell death rate on cancer cells, even at large tumor sizes, which can lead to cancer elimination. That would be the whole story, except that T cell killing rates can be diminished by exogenous factors in the tumor microenvironment. This necessarily requires a dynamical model capable of accounting for diversity in the microenvironmental influences on T cell killing rates, resulting from neighboring passive and actively immunoinhibitory cells (Fig. 1A).

**Fig. 1.**
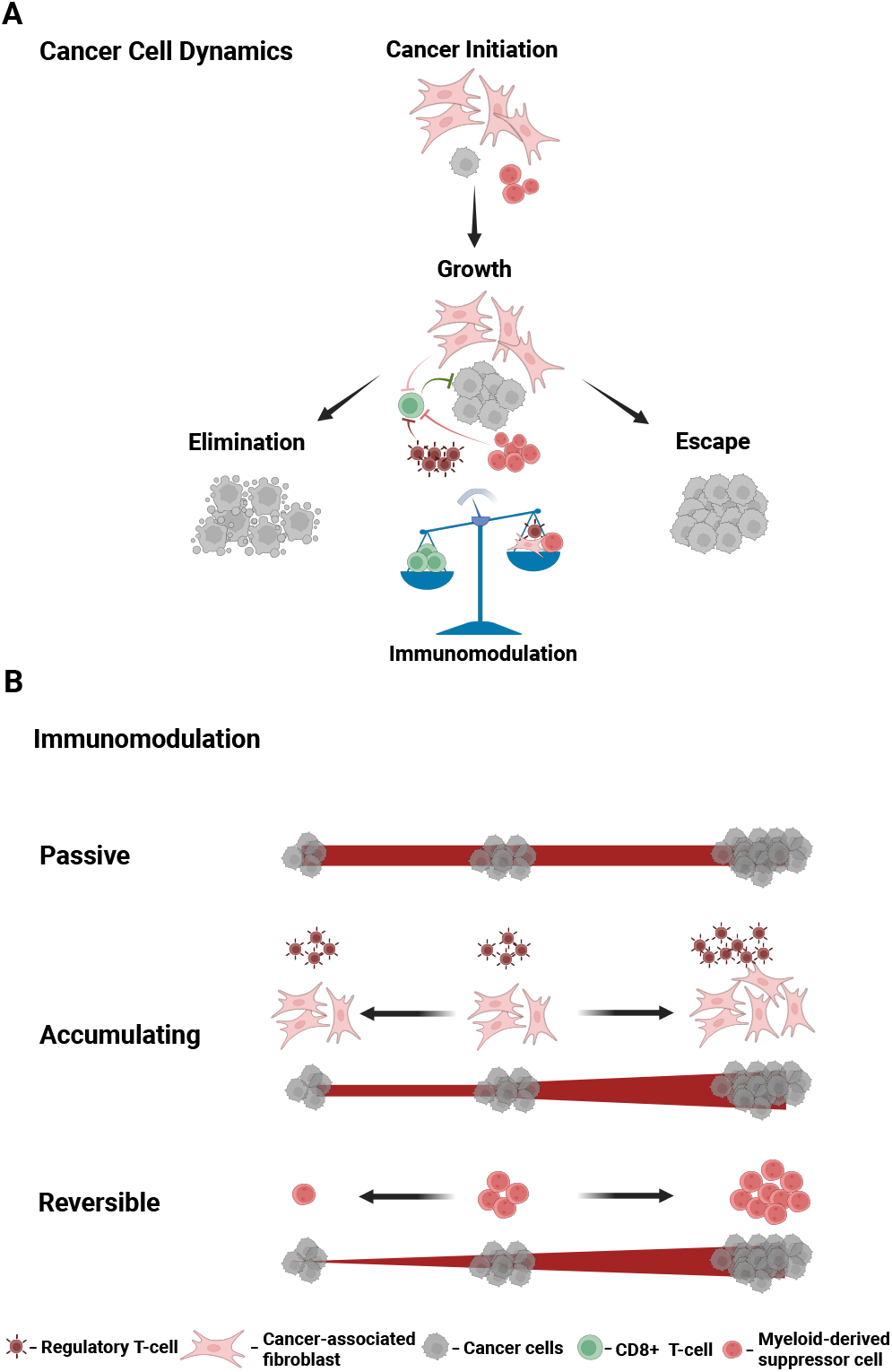
A) Conceptual schematic illustrating the tumor-immune dynamics that would lead to elimination or escape for the tumor population. The scale represents the relative balance between recognizing effector T cells, which eliminate the cancer, and the immunomodulatory factors that can diminish T cell killing and lead to immune escape. B) Immunomodulatory cells in the tumor microenvironment can act to impair T cell killing in a variety of ways. In the simplest “passive” case, the number of immunomodulatory cells is assumed to be fixed and independent from the number of cancer cells (*M* constant). Alternatively, the number of immunomodulatory cells may vary as a function of cancer cell number. In the “accumulating” case, the largest size that tumor population reaches determines *M* . This immunomodulation models the effects of Tregs and cancer-associated fibroblasts. In constrast, “reversible” immunomodulation describes the case in which the number of immunomodulatory cells may grow and shrink as a function of the number of cancer cells. This case may be more appropriate for modeling myeloid-derived suppressor cells. Created with BioRender.com.

We account for possible reductions in the T cell killing rate by introducing an *immunomodulation function f* that depends on tumor size *n* and the abundance of immunoinhibitory agents *M* . In general, we represent this as

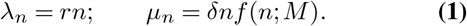

where *r, δ* with *δ* >> *r* are cancer per-capita growth and death rates, respectively. Here, *f* (*n*; *M* ) is the immunomodulation function. *f* ≤ 1 accounts for reductions in the T cell killing rate via *M* inhibitory cells. We consider both cases where *M* is constant or dynamic with respect to tumor size, and we refer to these as *static* and *dynamic immunomodulatory landscapes*, respectively. Refer to Section S8.2 of the SI for a general framework for the landscapes. We note that while there exist many reasonable choices for the precise functional form of *f* , for foundational understanding, we consider several specific cases below.

#### Static Immunomodulatory Landscape

In the simplest case, we assume that the tumor is surrounded by a fixed number of *M* neighboring stromal (nonmalignant) cells whose presence passively impairs T cell killing by diluting meaningful cancer-T cell interactions. We call this case *passive immunomodulation* with the functional form of *f* given by

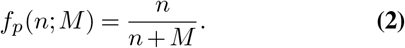

Here, *f* ≤ 1 reduces baseline T cell killing. We will show that T cell impairment via the mere presence of passive neighboring cells is incompatible with possible tumor escape. Furthermore, passive immunomodulation does not account for situations where the tumor microenvironment itself is immunosuppressive. For example, previous studies have suggested that T cell function is further impaired via metabolic dysfunction (38). Tumor cells together with glucose-depleted and lactate-rich environments can also lead to enhanced Treg suppressor activity (39).

We thus expand our model to consider *active immunomodulation*, wherein both the tumor cells and the neighbouring cells can modulate the level of T cell impairment. We list one such functional form wherein the death rate is inversely proportional to a linear combination of the inhibition signal and the tumor size. In this active case,

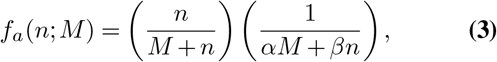

where *α* and *β* represent the effects induced by both types of cells in the tumor microenvironment.

#### Dynamic Immunomodulatory Landscape

Of course, it is also possible that immunomodulatory cells may vary dynamically with respect to the number of cancer cells. We also consider the more general scenario where changes in *M* are induced by changes in the tumor size. Motivated by cases where the growing tumor cells permanently recruit neighboring stromal cells (40) and also the reversible impact of MDSCs (35), we describe two distinct dynamical behaviors: one where the inhibitory signal, *M* , increases and decreases in response to the tumor population, and the other where immunoinhibitory signals depend on the cancer population size and, once present, exist permanently. We refer to these cases as *reversible* and *accumulating* immunomodulation, respectively (Fig 1B), and we calculate the hitting probabilities in these settings from an analytical perspective as well as a computationally tractable method.

### Diffusion Approximation

We construct a diffusion approximation of the birth-death process to further verify the results of the passive immunomodulation case with the understanding that the mathematical framework thus demonstrated applies to a larger class of birth-death processes. We show if the mean and variance expression for the birth-death model are Lipschitz, then one can find a diffusion approximation represented by a stochastic differential equation. We will establish a generalized approach for identifying a diffusion approximation represented by a stochastic differential equation for the broad class of birth-death processes whose mean and variance are Lipschitz. Using this approximation, we employ the full range of Itô calculus to obtain closed-form expressions for absorption probabilities as well as the mean absorption times.

### Numerical Simulation

All analytical results are compared to large-scale stochastic numerical simulation via the Gillespie algorithm.

### Data Analysis

Lastly, we apply these findings to cancer incidence data for patients from diagnosis to first progression, first progression to second progression, and so on to estimate the immunomodulation parameters from each event interval. In absence of clinical micrometastasis longitudinal tumor trajectory data, we characterize immune escape relative to individual progression events. We then calculate the distribution for progression time and subsequently fit our model. Best fit parameters were selected based on minimizing the mean squared error for the distribution of progression times for each progression event and for each cancer case. The breast cancer data was obtained from the website (41) associated with the breast cancer forecasting work by Newton et al. (42) using clinical data from Memorial Sloan Kettering Cancer Center and MD Anderson Cancer Center. Similarly, the bladder cancer data was obtained from the website (43) associated with the modelling work by Hasnain et al. (44) using clinical data from USC Institute of Urology.

## Results

In the following, we present the main findings of our analysis. Full mathematical derivations are provided in the Supplementary Information (SI). An overview is provided in SI Sec. S1.

### A. Passive Immunomodulation alone cannot promote cancer escape

If immune recognition is passively diminished by the presence of *M* neighboring nonmalignant cells as in Eq. (2), then the net growth rate, *ξ*(*x*), is given by

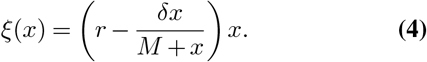

This behavior yields one unstable equilibrium point at the origin and one stable equilibrium point at *x*^*\**^ = *rM/*(*δ* − *r*) (Fig. 2A). Moreover, as a sequence of functions, the *ξ*(*x*) are increasing with respect to the inhibitory element *M*. The mean elimination time for this process, 𝔼[*T*_0_] (see SI Sec. S4), satisfies:

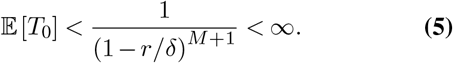

**Fig. 2.**
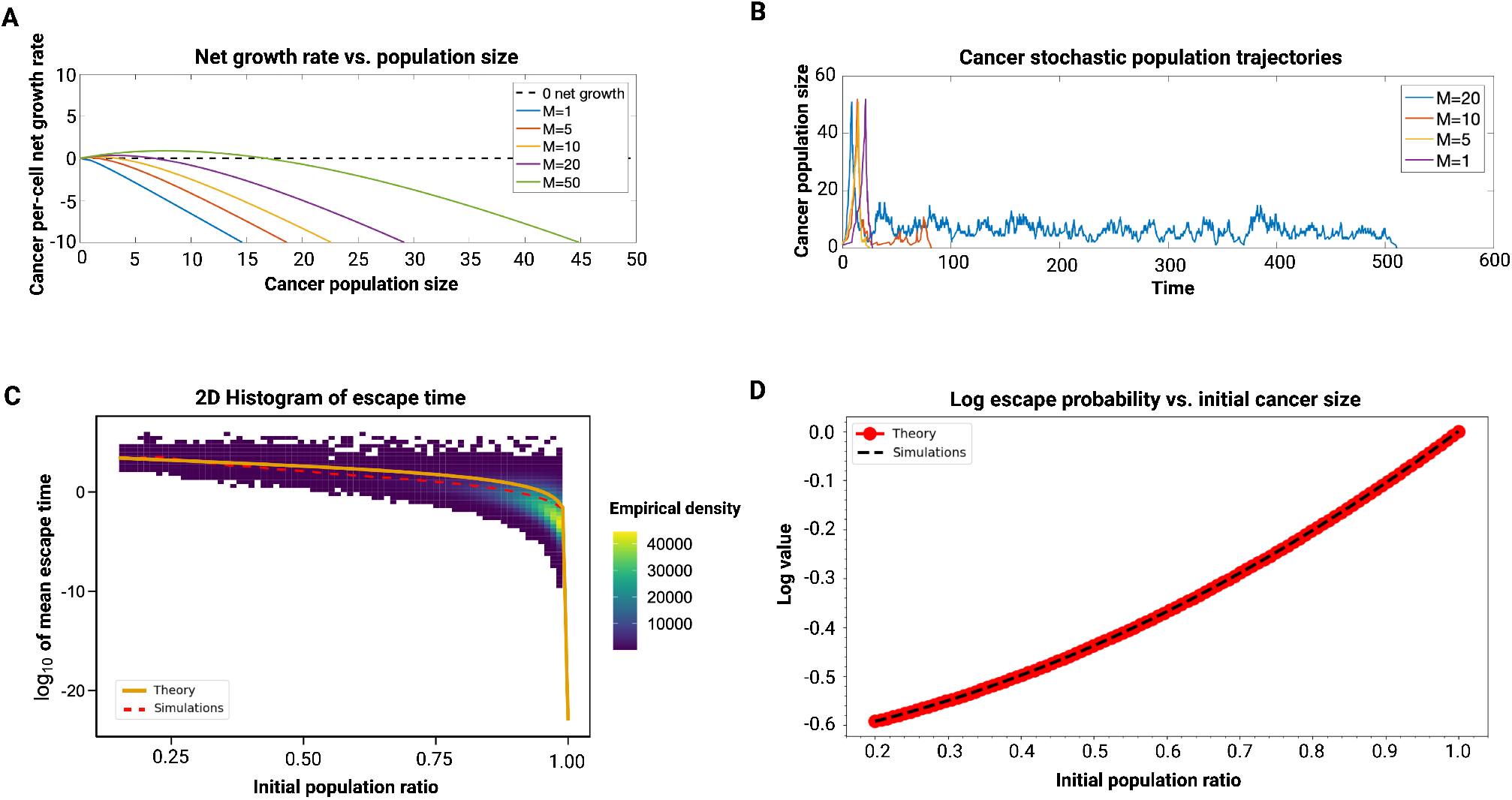
Dynamics under passive immunomodulation. A) Schematic of the cancer per-cell net growth rate as a function of cancer population size for various levels of immunomodulation (*r* = 0.25, *δ* = 1). B) Simulated stochastic trajectories with various immunomodulatory parameters (*r* = 0.25, *δ* = 1). Results of largescale stochastic simulations and analytical estimates for the diffusion approximation are provided for cancer population C) mean escape time and D) escape probability. The diffusion approximation for passive immunomodulation is given as a function of initial population ratio with *r* = 0.1, *δ* = 0.8, *N* = 100, and *M* = 2. Total simulations for initial population ratio under and over 0.25 are 100, 000 and 10, 000, respectively. Dashed lines represent simulated values and solid lines represent the analytical result obtained via diffusion approximation.

Collectively, these dynamics predict that under passive immunomodulation the population cannot grow to an arbitrarily large size but instead will fluctuate around *x*^*\**^ (Fig. 2B). However, the stochasticity of the birth-death process ultimately results in disease elimination after a finite average waiting time. *ξ*(*x*) > 0 for *x* < *x*^*\**^, which creates an barrier to elimination, and because *ξ* is an increasing function of *M* this barrier also increases as *M* grows, resulting in larger elimination times.

Under this model, appreciable disease only occurs if *x*^*\**^ ≫ 1, which, for effective T cell killing rates (*δ* − *r* > *r*), requires *M* to be many multiples larger than the size of macroscopically detectable disease. This finding, together with the absence of cancer escape, suggests passive immunomodulation alone is insufficient for explaining observed dynamics.

#### A.1. Diffusion approximation for general nonlinear birth-death processes

Despite its simplicity, the above case presents a prototypical example of a nonlinear birthdeath model wherein we desire statistical characterizations of escape for processes that can grow to large sizes prior to reaching an absorption state. Motivated by a desire to develop analytically tractable estimates of elimination probabilities and mean escape times for a variety of birthdeath processes, we next developed a general diffusion approximation-based approach.

Consider a birth-death process *X*^*N*^ (*t*) on a state space {0, 1, *…* , *N*} for *N* ≫ 1 representing the absorption size at cancer escape, with transition rates given by Eq. (1). Then we can construct an approximation *Z*^*N*^ (*t*) defined by the stochastic differential equation:

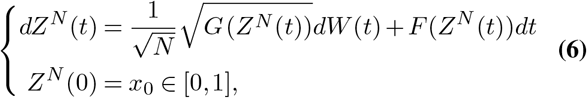

where *W* (*t*) is a standard Brownian motion, *F* (*x*) = *λ*_*x* −_ *µ*_*x*_ and *G*(*x*) = *λ*_*x*_ + *µ*_*x*_. Note that, *Z* = *X/N* tracks the population ratio with *Z* = 1 corresponding to cancer escape. We calculate the closed-form solution for ultimate extinction probability and time to escape given the initial population ratio *x*_0_. Let *U*_0_(*x*_0_) to be the probability that the diffusion process is ultimately eliminated. It can be shown (SI Section S6.2) that,

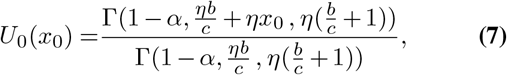

where *α, b, η, c* constants depend on *N, M, δ* , *r*, and Γ is the incomplete upper gamma function, given by

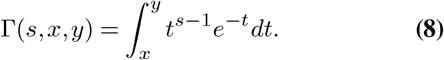

Similarly, if *τ* = inf {*t* > 0|*X*(*t*) = 1}, then the mean time for escape (SI Section S6.3) can be shown the by given by

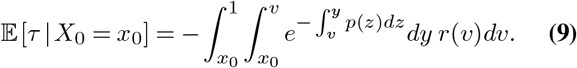

Both of these closed-form equations are exact for the diffusion approximation. Our approximation well-represents the birth-death process across all initial population fractions whenever escape is assumed to occur at a large population size (in general, *N ∼* 10^9^), as observed in comparing the analytical escape probability and mean time with those results obtained through largescale stochastic simulation (Fig. 2CD). We remark that this approach can be applied identically to study a variety of nonlinear birth-death processes, with the general requirement that *λ*_*x*_ and *µ*_*x*_ are Lipschitz in population fraction to guarantee asymptotic agreement between the process and its diffusion approximation for large *N* . The passive case, while it allows for long periods of non-extinction, precludes any reasonable possibility for tumor immune escape. Collectively, these findings motivate the analysis of a more realistic model of immunomodulation’s role in impairing T cell recognition.

### B. Active immunomodulation permits both tumor elimination and escape

In the more general case, active immunomodulatory behavior may be modeled by accounting for additional impairments in T cell function that result from increases in the inhibitory signal *M* or total tumor cell size *n*. Such immunomodulation more accurately describes T cell dysfunction resulting from, for example, resource consumption (e.g., depletion of L-arginine or cystine), which can reduce T cell killing efficiency directly and increase Treg suppressor activity (38, 39, 45). Such impairments be accounted for by incorporating an additional term in the immunomodulation function that varies inversely with respect to cancer and immunomodulatory cell numbers. Toward this end, we considered a variety of possible functional forms (SI Sec. S7) and focus our main discussion here on one particular case that possessed representative dynamics. In this case, an additional (*αM* + *βn*)^−1^ term is added to the passive immunomodulation function Eq. (3) to account for reductions in the cancer per-capita T cell killing rates as a function of total cell abundance.

In systems employing the general dynamics given by Eq. 1, the asymptotic behavior *f*_*∞*_ = lim_*n*→*∞*_ *f* (*n*) of the immunomodulatory function *f* can be partitioned by *r/δ*, which influences ultimate escape dynamics through the stability of the leading equilibrium point (See SI Sec. S5 for full details). Control occurs if *f*_*∞*_ < *r/δ*, while ultimate escape to large sizes occurs if *f*_*∞*_ > *r/δ*. In particular, passive immunomodulation yields *f*_*∞*_ = 1 > *r/δ*, resulting in stable cancer control, while active immunomodulation gives *f*_*∞*_ < *r/δ*, permitting cancer immune escape.

As such, the active immunomodulatory function *f*_*a*_ yields the size-inhomogeneous net-growth rate:

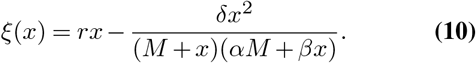

The nontrivial equilibrium points can be found by equating Eq. (10) to zero and solving

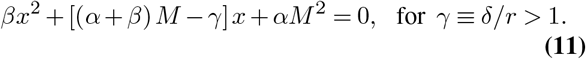

Since the above equation is quadratic in *x*, equilibrium points and their stability depend on the discriminant, *D*(*α, β*) of Eq. (11):

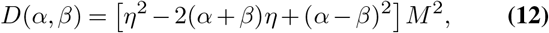

for *η* ≡ *δ/rM* , where *α* (rsp. *β*) controls the strength of inhibition for every present inhibitory element (resp. cancer cell). If we re-normalize with respect to *β*, Eq. (11), and hence equilibria of Eq. (10), can be fully characterized by *α*. This affords a variety of variety of dynamical behavior (SI Sec. S7.2). The most interesting cases possess both stable and unstable equilibrium cell sizes (SI Fig. S3). We focus our subsequent analysis to this case.

Fig 3 illustrate the behavior for the cancer net growth rate for various choices of the immunomodulatory parameter, *M* . The population regime of net-positive growth separating the origin from the first nonzero stable equilibrium state serves as a barrier to cancer elimination. Similarly, the population regime of net-negative growth rates above the first stable equilibrium population size to the population size of the nonstable equilibrium point represents a barrier to immune escape. Here, the stable and unstable states are separated by an activation barrier below which the population can be controlled by T cell recognition, and above which the microenvironment represses T cell killing enough to facilitate cancer escape. Hence a population existing at a stable equilibrium state would traverse either the elimination barrier to the left and become extinct, or it would cross the activation barrier to the right and undergo immune escape.

**Fig. 3.**
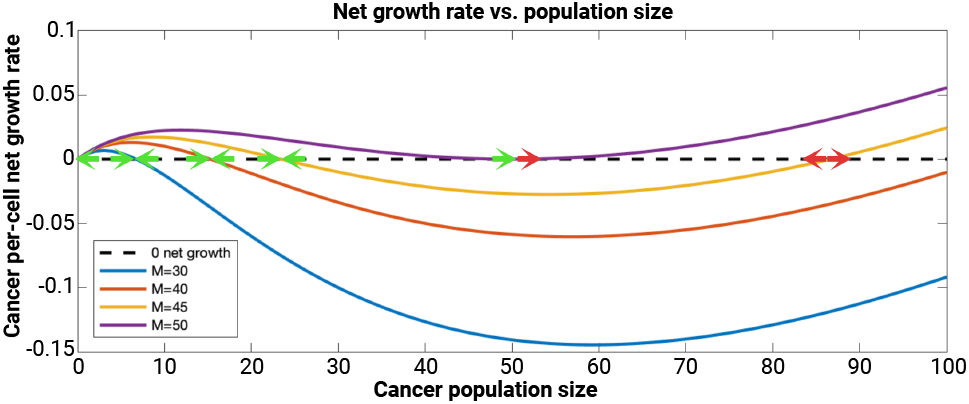
Schematic for the cancer net growth rates under active immunomodulation. Increases in immune inhibition in the tumor microenvironment (via increases in *M* ) lead to increases in the elimination barrier (values of the net growth curve from population size *n* = 1 until the first positive population size having zero net growth), and reduced barriers to immune escape (negative values of the net growth curve between the largest stable equilibrium point and detection size). In all cases, double green arrows indicate stable equilibrium points, double red arrows indicate unstable equilibrium points, and one of each indicates a semistable equilibrium point.

We remark that *ξ* is an increasing function of *M* , and so larger immunomodulaton leads to to larger equilibrium population sizes, larger elimination barriers, and reduced escape barriers. These dynamics allow a population to oscillate around the stable equilibrium for, perhaps large waiting times, prior to ultimate escape or elimination, and as a result more adequately model tumor-immune dynamics (21). Additionally, this more general framework resembles passive immunomodulation in the regime of small *M* , where the leading unstable equilibrium, in general, can become comparable to the upper detection size, effectively preventing immune escape.

### C. Dynamic Immunomodulatory Inhibition

Until now our model has assumed that the inhibitory signal, *M* , was constant throughout the evolution of the stochastic process. In reality, fluctuating cancer population sizes may also drive dynamic changes in immunomodulation present in the tumor microenvironment. For example, tumor-derived cytokines lead to Treg accumulation and can persist independent of tumor reduction (28). Similarly, cancer-associated fibroblasts that persist even after large reductions in tumor size can promote cancer cell survival (29). These examples would therefore be modeled more accurately assuming that immunomodulation *accumulates* as a tumor population grows, allowing for permanent recruitment of additional cells which remain the the tumor microenvironment irrespective of any future reductions in cancer size. Such a model contrasts with the behavior of myeloid-derived suppressor cells (MDSCs), another cell type capable of T cell impairment (32). In this case, MDSCs are present in the tumor microenvironment, and empirical evidence has demonstrated that tumor removal leads to MDSC reduction, suggesting a *reversible* mechanism of immunomodulation (35) (See SI Sec. S2.1 for additional details). Motivated by these biological phenomena, we provide a further generalization of the above modeling framework capable of accounting for immunomodulatory parameters that vary dynamically as a function of the tumor size. To do so, we characterize the hitting and absorption probabilities.

#### C.1. Conditional Hitting Probabilities

Escape probability calculations are more involved in this case, since the level of immunomodulation at fixed population sizes now also depends on the dynamic variable *M* . To characterize immune escape probability, we first consider the probability that the cancer population reaches a particular size *n* starting from an initial size *m* conditional on not hitting size *d*, denoted by event ^*d*^*E*_*n*,*m*_, for *d* < *m* < *n*. We first take an analytical approach to show that, for *n* ≥ *d* + 1,

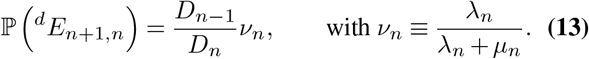

The *D*_*n*_ in Eq. (13) is defined as a solution of the recursive relation, given by

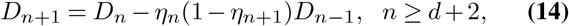

with *D*_*d*_ = *D*_*d*+1_ = 1. We identify the exact solution to this recursion relation (SI Sec. S8), however its implementation is computationally inefficient for large population sizes. For ease of calculations, we recast the recursion problem as a linear fractional transformation from real numbers to real numbers to evaluate the hitting probability by viewing it as a product of matrices. Namely,

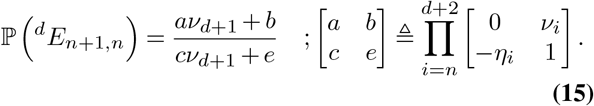

To apply this machinery to our problem, we characterize the dynamics using two sequences: the first sequence {*δ*_1_, *δ*_2_, … }, *δ*_*i*_ ≥ 0 is a collection of cancer population sizes at which the value of *M* changes. {*M*_1_, *M*_2_, … }, *M*_*i*_ ≥ 0 represents the corresponding immunomodulation values, so that *M* = *M*_*i*_ whenever *δ*_*i*_ ≤ *X* < *δ*_*i*+1_. The birth and death rates used to calculate the relevant conditional probabilities are functions of both *n* and *M* , given by *λ*_*n*,*M*_ and *µ*_*n*,*M*_ , respectively. We assume that the cancer population starts at initial size *n*_*i*_ with *M*_*i*_ suppressors. We remark that this framework is sufficiently general in its ability to handle any immune suppressor cell dynamics that can be represented by arbitrary {*δ*_*i*_} and corresponding {*M*_*i*_} sequences.

#### C.2. Accumulating Landscape

In this case, immunoinhibition *M* grows larger with increasing cancer population size and, once present, exists permanently. Thus, *M* (*t*) increasing in time at ever point that a new certain cancer population size threshold is achieved in the *δ* sequence. *M* is thus related to the maximal threshold size that the cancer population has currently achieved:

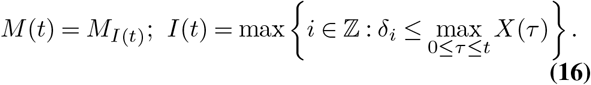

Using this formulation, the probability of immune escape can be calculated as a function of “step-up” probabilities, which are the conditional hitting probabilities between adjacent values of the *δ* sequence. Namely,

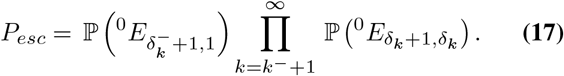

Using this approach, we calculate and compare the escape probabilities for the accumulating, reversible and the constant landscape using Eq. (13) and Eq. (15) (Fig. 4 and Fig. 5 respesctively). Our results demonstrate that an accumulating landscape pushes the tumor population to explosion as the ability to irreversibly recruit more inhibitory cells further catalyzes cancer escape at larger sizes. The ultimate effect is a reduced escape barrier and more likely escape for comparable initial values for *M* .

**Fig. 4.**
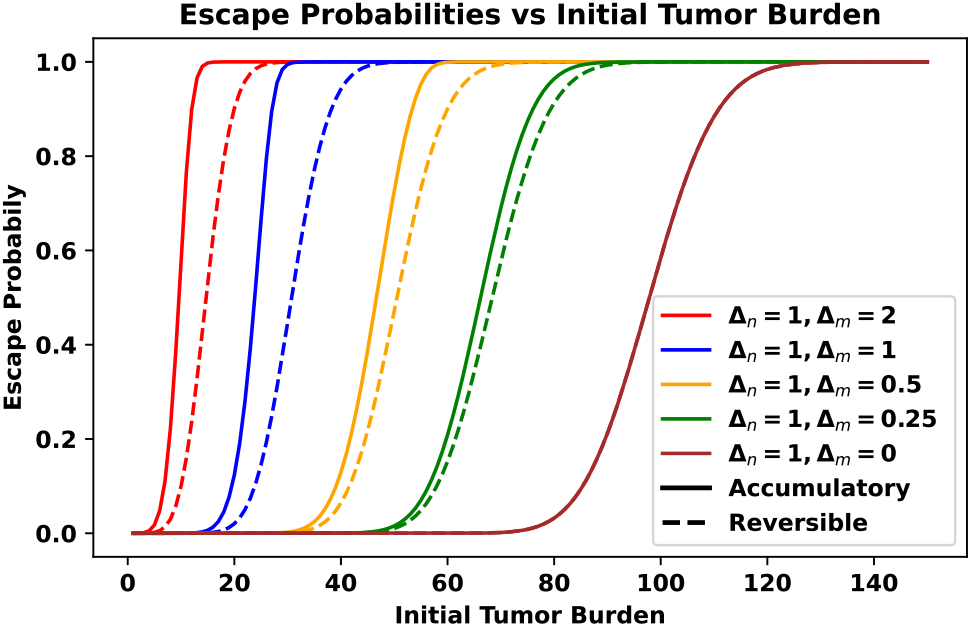
Escape Probabilities obtained analytically under dynamic immunomodulatory landscapes. Every Δ_*n*_ change (increase for accumulating) in tumor size results in Δ_*M*_ change (increase resp.) in the immunomodulatory signal. The brown line (Δ_*n*_ = 1, Δ_*M*_ = 0) is a special case that represents the escape probability under constant immunomodulation. In all cases, *f* is used according to linear inhibition case in Eq. (3) with *r* = 0.05, *δ* = 0.9, *N* = 150, *α* = 0.1, *M*_0_ = 1 , *β* = 0.18.

**Fig. 5.**
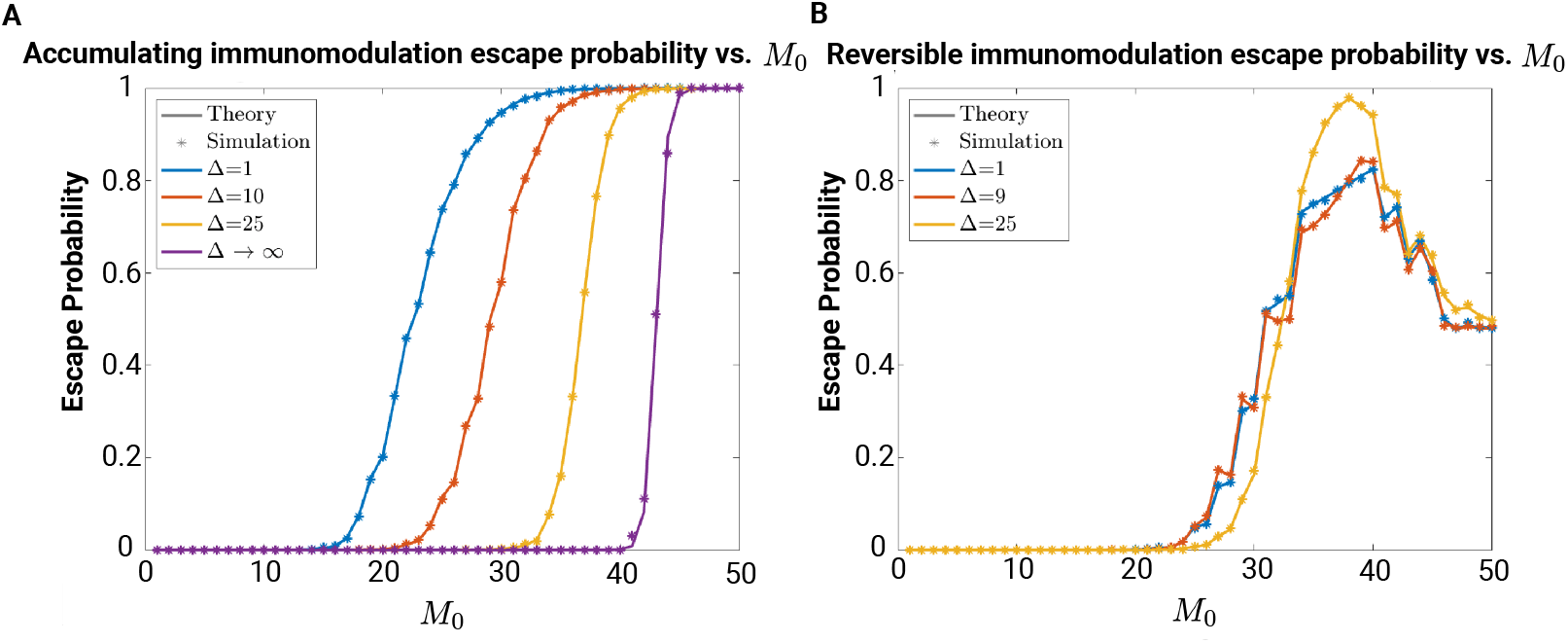
Escape probability under accumulating and reversible immunomodulation landscapes. Simulation estimates of escape probability are compared alongside the theoretical results (SI Sec. S8.1) for (A) accumulating and (B) reversible immunomodulation. For each curve, a total of 10^4^ stochastic simulations were used to obtain an average (in all cases, *f* is used according to Eq. (3) with *r* = 0.005, *δ* = 1, *α* = *β* = 1).

#### C.3. Reversible Landscape

In contrast with accumulating immunomodulation, reversible immunomodulation allows for *M* to both increase and decrease in response to cancer population growth and reduction, respectively. In this case, *M* is thus related to the current threshold size that the cancer has currently achieved in the *δ* sequence:

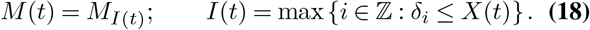

Largescale stochastic simulations for reversible and accumulating landscapes under a variety of step sizes Δ = *δ*_*i*+1_ − *δ*_*i*_ were compared to our analytical calculations of escape probability, indicating excellent agreement (Fig. 5 and Figs. S7, S8, and S9 in the SI). Intriguingly, the reversible case experiences nonmonotonic behavior in cancer escape owing to the fact that *M* may reversibly decline during changes in tumor size. From the linear fractional transform approach using Eq. (15) (Fig. 4), our results demonstrate that the reversible landscape offers similar benefits to a growing tumor as the accumulating case thus also catalyzing cancer escape at larger sizes. However, the reversible decline in the immodulatory signal during tumor progression leads to a later escape as compared to the accumulating case. We achieve agreement between calculated and simulated values for both reversible and accumulating immunomodulation landscapes across a range of partition intervals Δ = *δ*_*i*+1_ − *δ*_*i*_ and initial immunomodulation values *M*_0_ (Fig. 5).

### D. Application to Clinical Data

The above nonlinear birth-death processes illustrate how the particular mechanism underlying T cell impairment can lead to variable population dynamical responses. Given this, we next aimed to apply our framework to large-cohort studies in order to quantify which frameworks were more applicable for explaining observed distributions of cancer progression times, in addition to predicting the corresponding extent of immune impairment. We reasoned that the application of our model to subsequent cancer progression events would provide one way of assessing immune impairment underlying repeated disease escape, given the absence of detailed time-course descriptions tracking the population sizes of non-malignant cells in a small tumor microenvironment. Two excellent patient cohorts exist that contain information on time from diagnosis to first progression and subsequent inter-progression times.

These datasets comprise progression-free survival times for 3505 bladder cancer patients following diagnosis and cystectomy and 4181 breast cancer patients (41–44). In all cases, absence of metastatic disease was confirmed at the time of detection.

Toward that end, we calculated the empirical distributions for the *k*^th^ inter-progression times. These distributions were then compared to our theoretical results by fitting tumor escape times assuming dynamic (constant), reversible, and accumulating landscapes. The extent of immunomodulation *M* , along with *α* and *δ*, were used as free parameters for each inter-progression event (Fig. 6). As one might expect, reasonable fits for bladder and breast cases required distinct *α* and *δ* parameters for each cancer subtype, although in our fitting procedure we required that the optimized parameter choices be fixed for each family of progression curves within a given cancer subtype. We then identified the least-squares best-fit curves and corresponding *M* values while keeping the remaining dynamical parameters (*r* and *β*) fixed across all times for a given cancer type. In particular, the value of *M* was fitted in the static immunomodulation case, while in the active cases we fitted the fixed increment Δ*M* ≡ *M*_*i*+1_ − *M*_*i*_ having fixed initial value (*M*_0_ = 5). This procedure was applied assuming the active, reversible, and accumulating immunomodulation cases for both bladder and breast cancer (Fig. 6; MSE in SI Fig. S13).

**Fig. 6.**
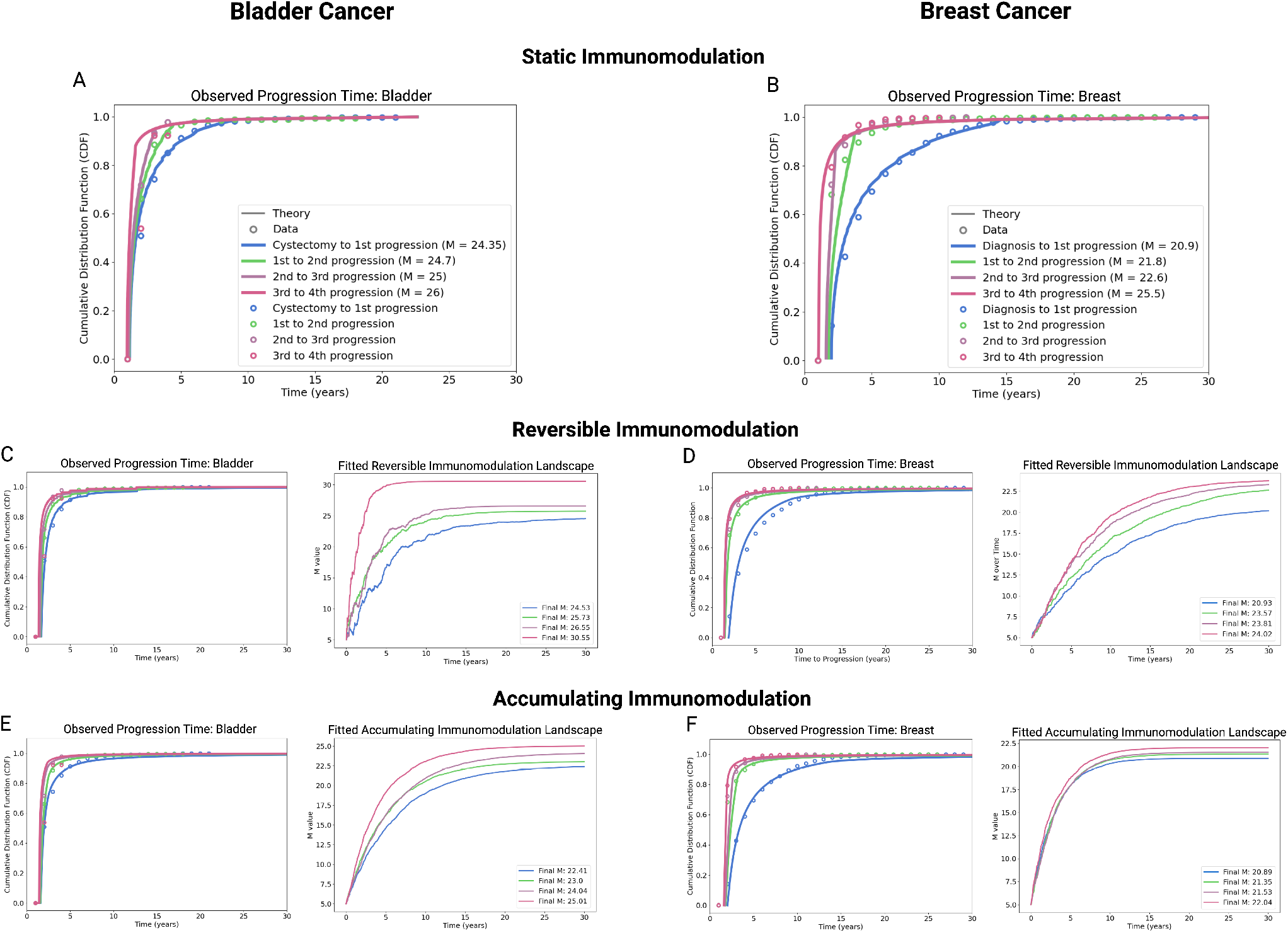
The mathematical model was applied to clinical data for bladder cancer (A) and breast cancer (B) to estimate the immunomodulatory parameter *M* under a constant immunomodulatory landscape. The parameters for all cases were set at *r* = 0.01 and *β* = 1. For bladder cancer, the fitted parameters were *α* = 0.81 and *δ* = 0.895, while for breast cancer, *α* = 0.81 and *δ* = 0.82. The model was also examined under a reversible immunomodulatory landscape, with *α* = 0.75 and *δ* = 0.99 for bladder cancer, and *α* = 0.85 and *δ* = 0.85 for breast cancer. Additionally, an accumulating immunomodulatory landscape was analyzed, as represented in Panels E and F. For bladder cancer (E), *α* = 0.93 and *δ* = 0.95, and for breast cancer (F), *α* = 0.85 and *δ* = 0.85. The initial value of *M* in both the accumulating and reversible cases was set to 5, with *M* incrementing by 0.01 in response to an increase of 10 cells in the population. These parameters, identified by minimizing the mean squared error (MSE) between the model’s predictions and clinical data, were determined to provide the best fit. Although the parameters differ between cancer types, they remain consistent across the stages of progression within each cancer type.

Our findings suggest that incorporation of the dynamic immunomodulation landscapes improves model fit for both breast and bladder cancer. Reversible immunomodulation outperforms accumulating immunomodulation in explaining observed progression for both breast and bladder cancer, although this finding is more pronounced in breast. Moreover, all three model descriptions appear to poorly explain advanced progression events in bladder cancer. Despite the reversible case being the best at explaining both disease types, the time evolution of *M* in the reversible case exhibits gradual increases in time which closely mimic the accumulating case. Within a given cancer subtype, the immunomodulatory signal strength increases from one progression step to the next. In comparing our results for a given model across cancer types, we find uniformly higher immune inhibitory values consistently occur in bladder cancer. This finding offers one plausible explanation for its relative aggressiveness from the standpoint of adaptive immune escape, which is consistent with observed immune escape in bladder cancer (46, 47)

In restricting our attention to reversible immunomodulation (Fig. 6C-D), both breast and bladder cancer are predicted to exhibit increasing immunomdulation landscapes as a function of progression events. We observe that the greatest jumps in the immunomodulation landscape are predicted to occur during early (first and second) and late (third and fourth) progression events for bladder cancer. This contrasts with breast cancer, which appear to demonstrate diminishing enhancements in immunomodulation over each subsequent progression event. In summary, our approach provides one method of mapping observed differences in relative cancer escape times to the underlying nature and extent of immune escape resulting from T cell recognition impairment.

## Discussion

Recent successes in cancer immunotherapy have delivered improved disease-free survival by enhancing the anti-tumor killing effect of T cells. However, it is known that the tumor-immune interaction may lead to cancer escape, elimination, or an equilibrium state wherein the immune system keeps cancer division under control. We developed a stochastic population dynamical model to account for both the role of immune impairment on cancer escape and also the nontrivial influence of fluctuations on cancer elimination in the regime of small population sizes. Our generalized theoretical framework, together with the diffusion approximation and quantification of the likelihood and timing of escape, is applicable to other biological contexts where populations undergo nonlinear birth and death.

In a model of balanced tumor growth and death resulting in stable cancer equilibrium states at small population sizes, ultimate cancer escape is infeasible if T cell impairment results passively from the mere presence of surrounding stromal cells. This led us to extend our model to account for active immunomodulation based on the population size of cancer and inhibitory cells. The extended model can flexibly describe a variety of active impairments, including in the setting where inhibitory cells vary dynamically in response to the number of (stochastically fluctuating) cancer cells. In our application to repeated cancer progression events, our model fits suggest the relative suitability of a reversible immune impairment in the tumor microenvironment for both breast and bladder cancer.

Of course, many additional factors, including treatment course, cancer heterogeneity, and individual patient characteristics, all play an important role in determining progression times. Our foundational model offers a minimalistic description of how varying immune impairment alone can result in disease progression, which already does a reasonable job in fitting families of these distributions. Variation in the extent to which each subsequent progression event altered the predicted immune inhibitory landscape provides one way where such a model can be employed to plan aggressive treatment vs. conservative management. The finding that a reversible immunomodulation framework best explains the empirical data bears potential implications for optimal treatment and also offers hope that the immune microenvironment can be “coaxed” into a state that favors cancer elimination. Such results are of clinical significance and require further experimental follow-up.

As mentioned above, one model strength is the relative simplicity of casting immune impairment as a nonlinear term in the cancer death rate. Such an approach, while amenable to analytic characterization, does not explicitly track the complex dynamics involving multiple cell types, signaling molecules, and feedback mechanisms, which are coarsegrained into the net growth parameter directly. In particular, the detailed dynamics of T cell activation, proliferation, and suppression are not explicitly modeled, nor are other immune populations tracked. Population dynamics are determined assuming homogeneous interactions between the cancer and immune cells, while in reality, spatial effects (such as T cell exclusion) may distort this idealized representation in solid malignancies. This model does not track distinct tumor and T cell clonotypes and consequent co-evolution process. Such an extension could be considered by assigning clone-specific parameter values, as has been the focus of prior studies (11, 13). We considered a dynamical immunomodulation parameter, but it is also possible that the other governing parameters are dynamic variables in time. Lastly, and though a relatively lenient assumption, the diffusion approximation is limited to describing Lipschitz birth and death rates (SI. Sec. S6). Despite these considerations, this work represents the first stochastic description and corresponding analytic theory of tumor immune-mediated dormancy. Subsequent work will be directed validating model predictions on experimental data and building into the model immunotherapeutic interventions to predict maximally effective interventions.

## ACKNOWLEDGEMENTS

We thank Herbert Levine for helpful discussions on modeling cancer immunemediated dormancy. JTG was supported by the Cancer Prevention Research Institute of Texas (RR210080). JTG is a CPRIT Scholar in Cancer Research.

## Bibliography

1. Franziska Michor, Yoh Iwasa, and Martin A Nowak. Dynamics of cancer progression. Nature reviews cancer, 4(3):197–205, 2004.

2. Yoh Iwasa, Martin A Nowak, and Franziska Michor. Evolution of resistance during clonal expansion. Genetics, 172(4):2557–2566, 2006.

3. Natalia Komarova. Stochastic modeling of drug resistance in cancer. Journal of theoretical biology, 239(3):351–366, 2006.

4. Aaron N Hata, Matthew J Niederst, Hannah L Archibald, Maria Gomez-Caraballo, Faria M Siddiqui, Hillary E Mulvey, Yosef E Maruvka, Fei Ji, Hyo-eun C Bhang, Viveksagar Krishnamurthy Radhakrishna, et al. Tumor cells can follow distinct evolutionary paths to become resistant to epidermal growth factor receptor inhibition. Nature medicine, 22(3):262–269, 2016.

5. Luis A Diaz Jr, Richard T Williams, Jian Wu, Isaac Kinde, J Randolph Hecht, Jordan Berlin, Benjamin Allen, Ivana Bozic, Johannes G Reiter, Martin A Nowak, et al. The molecular evolution of acquired resistance to targeted egfr blockade in colorectal cancers. Nature, 486(7404):537–540, 2012.

6. Jennifer Couzin-Frankel. Cancer immunotherapy. Science, 342(6165):1432–1433, 2013.

7. Theresa L Whiteside. Immune suppression in cancer: effects on immune cells, mechanisms and future therapeutic intervention. Seminars in cancer biology, 16(1):3–15, 2006.

8. Robert D Schreiber, Lloyd J Old, and Mark J Smyth. Cancer immunoediting: integrating immunity’s roles in cancer suppression and promotion. Science, 331(6024):1565–1570, 2011.

9. Jason T George and Herbert Levine. Stochastic modeling of tumor progression and immune evasion. Journal of theoretical biology, 458:148–155, 2018.

10. Sam Palmer, Luca Albergante, Clare C Blackburn, and TJ Newman. Thymic involution and rising disease incidence with age. Proceedings of the National Academy of Sciences, 115 (8):1883–1888, 2018.

11. Jason T George and Herbert Levine. Implications of tumor–immune coevolution on cancer evasion and optimized immunotherapy. Trends in Cancer, 7(4):P373–383, 2021.

12. Jason T George and Herbert Levine. Sustained coevolution in a stochastic model of cancer– immune interaction. Cancer research, 80(4):811–819, 2020.

13. Jason T George and Herbert Levine. Optimal cancer evasion in a dynamic immune microenvironment generates diverse post-escape tumor antigenicity profiles. Elife, 12:e82786, 2023.

14. Samra Turajlic, Hang Xu, Kevin Litchfield, Andrew Rowan, Tim Chambers, Jose I Lopez, David Nicol, Tim O’Brien, James Larkin, Stuart Horswell, et al. Tracking cancer evolution reveals constrained routes to metastases: Tracerx renal. Cell, 173(3):581–594, 2018.

15. Mariam Jamal-Hanjani, Gareth A Wilson, Nicholas McGranahan, Nicolai J Birkbak, Thomas BK Watkins, Selvaraju Veeriah, Seema Shafi, Diana H Johnson, Richard Mitter, Rachel Rosenthal, et al. Tracking the evolution of non–small-cell lung cancer. New England Journal of Medicine, 376(22):2109–2121, 2017.

16. Nicholas McGranahan and Charles Swanton. Clonal heterogeneity and tumor evolution: past, present, and the future. Cell, 168(4):613–628, 2017.

17. Eszter Lakatos, Marc J Williams, Ryan O Schenck, William CH Cross, Jacob Househam, Luis Zapata, Benjamin Werner, Chandler Gatenbee, Mark Robertson-Tessi, Chris P Barnes, et al. Evolutionary dynamics of neoantigens in growing tumors. Nature genetics, 52(10): 1057–1066, 2020.

18. Filippo G Giancotti. Mechanisms governing metastatic dormancy and reactivation. Cell, 155(4):750–764, 2013.

19. Albert C Yeh and Sridhar Ramaswamy. Mechanisms of cancer cell dormancy—another hallmark of cancer? Cancer research, 75(23):5014–5022, 2015.

20. Hao-fan Wang, Sha-sha Wang, Mei-chang Huang, Xin-hua Liang, Ya-Jie Tang, and Ya-ling Tang. Targeting immune-mediated dormancy: a promising treatment of cancer. Frontiers in oncology, 9:498, 2019.

21. Gavin P Dunn, Lloyd J Old, and Robert D Schreiber. The three es of cancer immunoediting. Annu. Rev. Immunol., 22:329–360, 2004.

22. Rona M MacKie, Robin Reid, and Brian Junor. Fatal melanoma transferred in a donated kidney 16 years after melanoma surgery. New England Journal of Medicine, 348(6):567– 568, 2003.

23. Catherine M Koebel, William Vermi, Jeremy B Swann, Nadeen Zerafa, Scott J Rodig, Lloyd J Old, Mark J Smyth, and Robert D Schreiber. Adaptive immunity maintains occult cancer in an equilibrium state. Nature, 450(7171):903–907, 2007.

24. Matthew A Summers, Michelle M McDonald, and Peter I Croucher. Cancer cell dormancy in metastasis. Cold Spring Harbor Perspectives in Medicine, 10(4), 2020.

25. Barbara Molon, Bianca Calì, and Antonella Viola. T cells and cancer: how metabolism shapes immunity. Frontiers in immunology, 7:20, 2016.

26. Amedeo Amedei, Elena Niccolai, Marisa Benagiano, Chiara Della Bella, Fabio Cianchi, Paolo Bechi, Antonio Taddei, Lapo Bencini, Marco Farsi, Paola Cappello, et al. Ex vivo analysis of pancreatic cancer-infiltrating t lymphocytes reveals that eno-specific tregs accumulate in tumor tissue and inhibit th1/th17 effector cell functions. Cancer Immunology, Immunotherapy, 62:1249–1260, 2013.

27. Xuefang Cao, Sheng F Cai, Todd A Fehniger, Jiling Song, Lynne I Collins, David R Piwnica-Worms, and Timothy J Ley. Granzyme b and perforin are important for regulatory t cell-mediated suppression of tumor clearance. Immunity, 27(4):635–646, 2007.

28. Theresa L Whiteside. The role of regulatory t cells in cancer immunology. ImmunoTargets and therapy, pages 159–171, 2015.

29. Ralf-Peter Czekay, Dong-Joo Cheon, Rohan Samarakoon, Stacie M Kutz, and Paul J Higgins. Cancer-associated fibroblasts: mechanisms of tumor progression and novel therapeutic targets. Cancers, 14(5):1231, 2022.

30. Raisa A Glabman, Peter L Choyke, and Noriko Sato. Cancer-associated fibroblasts: Tumorigenicity and targeting for cancer therapy. Cancers, 14(16):3906, 2022.

31. Aikaterini Hatziioannou, Themis Alissafi, and Panayotis Verginis. Myeloid-derived suppressor cells and t regulatory cells in tumors: unraveling the dark side of the force. Journal of Leukocyte Biology, 102(2):407–421, 2017.

32. Minu K Srivastava, Pratima Sinha, Virginia K Clements, Paulo Rodriguez, and Suzanne Ostrand-Rosenberg. Myeloid-derived suppressor cells inhibit t-cell activation by depleting cystine and cysteine. Cancer research, 70(1):68–77, 2010.

33. Ngozi R Monu and Alan B Frey. Myeloid-derived suppressor cells and anti-tumor t cells: a complex relationship. Immunological investigations, 41(6-7):595–613, 2012.

34. Bo Huang, Ping-Ying Pan, Qingsheng Li, Alice I Sato, David E Levy, Jonathan Bromberg, Celia M Divino, and Shu-Hsia Chen. Gr-1+ cd115+ immature myeloid suppressor cells mediate the development of tumor-induced t regulatory cells and t-cell anergy in tumor-bearing host. Cancer research, 66(2):1123–1131, 2006.

35. Cunren Liu, Shaohua Yu, John Kappes, Jianhua Wang, William E Grizzle, Kurt R Zinn, and Huang-Ge Zhang. Expansion of spleen myeloid suppressor cells represses nk cell cytotoxicity in tumor-bearing host. Blood, 109(10):4336–4342, 2007.

36. Kathleen P Wilkie and Philip Hahnfeldt. Mathematical models of immune-induced cancer dormancy and the emergence of immune evasion. Interface Focus, 3(4):20130010, 2013.

37. Kathleen P Wilkie, Philip Hahnfeldt, and Lynn Hlatky. Using ordinary differential equations to explore cancer-immune dynamics and tumor dormancy. bioRxiv, page 049874, 2016.

38. Nicole E Scharping, Ashley V Menk, Rebecca S Moreci, Ryan D Whetstone, Rebekah E Dadey, Simon C Watkins, Robert L Ferris, and Greg M Delgoffe. The tumor microenvironment represses t cell mitochondrial biogenesis to drive intratumoral t cell metabolic insufficiency and dysfunction. Immunity, 45(2):374–388, 2016.

39. McLane J Watson, Paolo DA Vignali, Steven J Mullett, Abigail E Overacre-Delgoffe, Ronal M Peralta, Stephanie Grebinoski, Ashley V Menk, Natalie L Rittenhouse, Kristin DePeaux, Ryan D Whetstone, et al. Metabolic support of tumour-infiltrating regulatory t cells by lactic acid. Nature, 591(7851):645–651, 2021.

40. K. C. Valkenburg, A. E. de Groot, and K. J. Pienta. Targeting the tumour stroma to improve cancer therapy. Nat Rev Clin Oncol, 15(6):366–381, Jun 2018.

41. Paul K. Newton, Jeremy Mason, Neethi Venkatappa, Maxine S. Jochelson, Brian Hurt, Jorge Nieva, Elizabeth Comen, Larry Norton, and Peter Kuhn. Breast cancer modelling, 2015. Website online available; accessed at 8/14/2024.

42. Paul K. Newton, Jeremy Mason, Neethi Venkatappa, Maxine S. Jochelson, Brian Hurt, Jorge Nieva, Elizabeth Comen, Larry Norton, and Peter Kuhn. Spatiotemporal progression of metastatic breast cancer: a markov chain model highlighting the role of early metastatic sites. npj Breast Cancer, 1(1):15018, Oct 2015. ISSN 2374-4677. doi: 10.1038/npjbcancer.2015.18.

43. Zaki Hasnain, Jeremy Mason, Karanvir Gill, Gus Miranda, Inderbir S. Gill, Peter Kuhn, and Paul K. Newton. Bladder cancer modelling, 2019. Website online available; accessed at 8/14/2024.

44. Zaki Hasnain, Jeremy Mason, Karanvir Gill, Gus Miranda, Inderbir S. Gill, Peter Kuhn, and Paul K. Newton. Machine learning models for predicting post-cystectomy recurrence and survival in bladder cancer patients. PLOS ONE, 14(2):1–15, 02 2019. doi: 10.1371/journal.pone.0210976.

45. Zhaonian Hao, Ruyuan Li, Yuanyuan Wang, Shuangying Li, Zhenya Hong, and Zhiqiang Han. Landscape of myeloid-derived suppressor cell in tumor immunotherapy. Biomarker Research, 9(1):1–28, 2021.

46. Hernani Gil-Julio, Francisco Perea, Antonio Rodriguez-Nicolas, Jose Manuel Cozar, Amanda Rocío González-Ramirez, Angel Concha, Federico Garrido, Natalia Aptsiauri, and Francisco Ruiz-Cabello. Tumor escape phenotype in bladder cancer is associated with loss of hla class i expression, t-cell exclusion and stromal changes. International Journal of Molecular Sciences, 22(14):7248, 2021.

47. Zhao Yang, Yinyan Xu, Ying Bi, Nan Zhang, Haifeng Wang, Tianying Xing, Suhang Bai, Zongyi Shen, Faiza Naz, Zichen Zhang, et al. Immune escape mechanisms and immunotherapy of urothelial bladder cancer. Journal of clinical and translational research, 7(4):485, 2021.

