## Supplementary Information for "How Modulation of the Tumor Microenvironment Drives Cancer Immune Escape Dynamics"

August 16, 2024

#### S1 Overview

Here we develop a generalized framework for studying the influence of stochastic immunomodulation on the growth dynamics of a cancer population. Sec. S2 describes an overview of the clinical problem and relevant cellular influences on immune killing. Sec. S3 frames the general mathematical problem of interest. Sec. S4 discusses the dynamics assuming that immunomodulation imparts a passive inhibition on T cell recognition in proportion to the number of immunomodulatory cells, with a conclusion that this method of impairment always leads to population control. Sec. S5 establishes sufficient conditions for equilibrium and the long-term behavior of escape or stable control. Sec. S6 develops a versatile diffusion approximation framework applicable to nonlinear birth-death process of the general form in Sec. S3, and establishes analytical elimination probabilities and mean cancer population escape times. Contrasting with passive inhibition, Sec. S7 next considers the effects of active immunomodulatory inhibition on T cell killing. Lastly, Sec. S8 extends the model to the physiologic case wherein immunomodulation depends on immunosuppressive cells that are dynamically recruited to the tumor microenvironment based on dynamical changes in cancer cell size.

### S2 Background and Motivation: Dormancy and Immunomodulation

Cancer dormancy, initially in reference to cancer cells in a state of mitotic quiescence [1] refers to the general belief that cancer cells may persist, despite possible therapeutic intervention, in a quiescent state. Since then, researchers have elaborated on possible physiological dormant states, including cellular dormancy and immune-mediated dormancy, with such states imparting chemotherapeutic resistance [2]. The competing roles pro-tumor and anti-tumor immunity and their influence on cancer elimination and progression has now been well-characterized. In this context, researchers have conceptualized this balance as possibly resulting in cancer escape, elimination, or equilibrium [3, 4]. In this context, immune-mediated dormancy is associated with the equilibrium phase, where the understanding is that T cell killing matches the cancer division rate [5].

Compelling experimental and clinical evidence exists in support of immune-mediated dormancy. In one striking example, two kidney transplant recipients sharing the same donor placed on chronic immunosuppression developed fatal melanoma [6]. In this case, the donor had undergone surgical resection of primary melanoma 16 years prior to kidney donation. In one recipient, primary breast cancer was diagnosed, prompting additional studies that revealed two subcutaneous nodules which were identified as secondary melanoma. In the other recipient, a palpable kidney lump prompted biopsy showing secondary melanoma. This case speaks to the immune dormancy of the donor disease, presumably awakening in the new setting of immunosuppression. One central working hypothesis is that early cancer dissemination produces dormant cells, which may later reemerge as metastatic disease [7]. In separate studies, carcinogen-injected mice that did not develop progressive malignancy, many of which having stable masses at the injection site, were treated with either control antibody or antibody depleting CD4+ and CD8+ T cells [8]. Those treated with immune-depleting factors developed progressive malignancy, in contrast to the control group. The experiment was repeated with mice lacking adaptive immunity and resulted in very few late-forming tumors, implicating the role of adaptive immunity in late tumor growth. Carcinogen injection site histology demonstrated atypia and vimentin expression in immunocompetent mice with delayed presentation. Immunocompetent mice with early presentation showed inflammation and fibrosis with multinucleated giant cells. Moreover, immune-depleted mice lacked atypical cells and giant cells. This study used the term 'equilibrium' to describe the stable tumor masses in a subpopulation of immunocompetent mice without progressive disease, suggesting the role of balanced tumor growth and immune killing.

An additional study focused specifically on the role of MHC-I expression levels in immune-mediated escape and elimination [9]. Metastatic dormancy, defined by 24 months free of metastasis following tumor removal, occurred in mice with intact adaptive immune systems. MHC-I expression was previously noted to be inversely correlated with tumor-initiating capacity *in vivo*. In this study, primary tumors grew rapidly, presumably due to baseline low MHC-I phenotypes. The authors noted a reversed trend in metastasis, such that MHC-I expression correlated directly with metastatic potential. Restoration of host immune responses via immunotherapy eradicated metastases. This led to a subsequent review arguing the necessity of MHC-I recovery in the setting of immunotherapy [10]. Here, the authors conceptualized escape, elimination, and dormancy around 3 cell phenotypes: Tumor MHC-I strongly positive cells, Escaped tumor cells MHC class I negative cells, and Dormant/awake weak to intermediate MHC class I cells. Interestingly, they suggested that the second category arises from the third, so that directionality exists in these states [10].

This body of work has lead to the general belief that the immune system keeps a population of tumor cells in check [11]. At the same time, (innate) immune-mediated inflammatory mechanisms may 'awaken' dormant cancer cells into aggressive metastases [12, 13]. Given our current understanding on tumor dormancy, there exists a significant need to resolve the scenario where cancer division is absent, termed 'solitary tumor cell dormancy' from the case where a growing population of cancer cells is kept in check by the immune system [11], termed 'micrometastatic dormancy', perhaps modulated by an environment that depends on the cancer population. Prior mathematical models have been proposed to study dormancy, including ODE models of cancer/immune system interactions [14,15]. These models, while capable of describing mean behavior immune-mediated dormancy, characterize cell sizes on a continuum. Moreover, predictions having equilibrium population sizes that are close to zero neglect the nontrivial extinction probability of this absorbing state. Their inability to resolve the distributional behavior of zero net-growth rates with positive division make them ill-equipped to differentiate between solitary and micrometastatic dormancy. Lastly, the recruitment of cellular immunomodulatory elements and their effect on T cell killing is also appreciated (discussed in Sec. S2.1 below), and their effects are complicated by the fact that arrivals are often dynamically coupled to the tumor population size. Given this, we develop a stochastic modeling framework to provide a more accurate distributional characterization of small cancer population dynamics, in addition to a dynamical description of immune impairment and cancer-mediated recruitment of inhibitory elements.

#### S2.1 Mechanisms of Immune Killing Impairment

Immune microenvironment-mediated inhibition can have a variety of effects on the T cell killing rate. We detail some of these below:

**T regulatory cells (Tregs):** Tregs are the major cellular components that regulate T cell killing activity, and therefore play a role in T cell-mediated cancer immune escape. Tregs play an important physiologic role in peripheral tolerance homeostasis. In cancer, these cells are found in increased abundance and often limit therapeutic intervention [16]. Tregs have been shown to limit the proliferation of cytotoxic T cells upon antigen-specific encounters, [17], in addition to Treg-specific granzyme B and perforin killing of CD8+ T cells [18]. Tumor-derived cytokines lead to Treg accumulation [19], and can persist independent of tumor reduction.

**Cancer-Associated Fibroblasts (CAFs):** In solid cancers, CAFs represent a phenotypically diverse group of mesenchymal cells. Their pro-tumorigenic presence has been associated with more aggressive disease, and they facilitate immune escape through various means, including direct inhibition of effector T cells and recruitment of Tregs [20, 21]. CAFs arise from resting fibroblasts, which are converted to an activated CAF state by cytokines present in the cancer and surrounding stroma, and they persist throughout tumor progression.

**Myeloid-derived suppressor cells (MDSCs):** Early experimental studies have detailed the dependence on T cells, which inherently lack cystathionase and hence an ability to synthesize cysteine, on exogenous cysteine provided by antigen-presenting cells [22]. MDSCs, also dependent on exogeneous cysteine, have been shown to sequester cysteine and by doing so impair T cell proliferation [23]. Additional impairments to the effector phase via failure to recruit tyrosine kinases in the T cell receptor (TCR) signaling pathway [24]. Lastly, MDSCs may themselves contribute to the presence of Tregs [25]. In contrast with Tregs and CAFs, empirical evidence has demonstrated that tumor removal leads to MDSC reduction, suggesting a reversible-mechanism of immunomodulation [26].

We restrict our attention to the case where the cancer population may effectively be targeted by cytotoxic T cells via recognition of tumor-specific antigenic signatures. The dynamics described below elaborate on cases where such population-level T cell targeting may be affected by exogeneous dependencies in the immune micro-environment. We neglect the effects of cancer-specific evolution resulting in escape by altering the set of possible T cell targets, for which we refer to a previous analysis on those general dynamics [27–29].

#### S3 Model Development

We will develop a general modeling framework by using time homogeneous non-linear birth-death processes to describe the tumor growth and killing behavior. Here, a cancer population of $N(t)$ cells undergoes time homogeneous stochastic birth and death. We will for foundational understanding further assume that the cancer population is driven by exponential growth for the sizes at which we are interested in studying with per-cell rate $r$ and that an active immune response mounted against a cancer population will induce a (large) per-cell death rate $\delta$, with possible modification by an immunomodulation function, $f(n;M)$, that may depend on the population size, $n$, and is modified by an inhibitory element $M$. Here, $M$ represents in abstraction an inhibitory signal on T cell killing. $f(n;M)$ is introduced as a general term that may tangibly represent, for example, inhibition due for example to a particular cell abundance or metabolic signal. Such application will determine differences in $f$, $M$, and $M's$ dynamics relative to $N(t)$. In this framework, the birth rate at size $n$, $\lambda_n$, and corresponding death rate, $\mu_n$ are given by

$$\lambda_n = rn; \qquad \mu_n = \delta n f(n;M). \tag{S1}$$

Previous models [27, 30] focused on the dynamics induced by an evading phenotype assumed killing at the maximal rate ($\mu_n = \delta n$) once the population size or growth-rate exceeded a lower detection threshold. This effectively eliminated all threats unable to acquire an immune evasive phenotype with per-cell division rates $r < \delta$. Here, we are interested in characterizing the effects of the microenvironment on T cell killing and cancer escape. We start by illustrating the simplest type of model in Sec. S4. This is followed in Sec. S5 by a discussion of the general model behavior, and then by additional relevant cases.

#### S4 Passive immunomodulatory participation

The simplest framework for our model assumes that the inhibitory element, $M$, is fixed and that immunomodulation is given by the abundance of cancer cells relative to inhibitory elements:

$$f(n; M) = \left(\frac{n}{M+n}\right). \tag{S2}$$

Tangibly, this framework models a collection of $M$ *passive cells*, such as stromal or other neighboring cell types. Their presence do not directly inhibit T cell killing, but instead impede detection indirectly by dilution of the T cell-cancer interaction simply based on their relative abundance. In this case the net growth rate $\xi(x) = \lambda_x - \mu_x$ is characterized by

$$\xi(x) = \left(r - \frac{\delta x}{M+x}\right)x \quad \text{and} \quad \xi'(x) = r - \frac{\delta(2M-x)x}{(M+x)^2}, \tag{S3}$$

which yield 2 equilibria:

$$x_1^* = 0, \qquad x_2^* = \frac{rM}{\delta - r}.$$

Here, $x_1^*$ is an absorbing state and it is unstable since $\lambda_n > \mu_n$ when $n < x_2^*$. Since $\lambda_n < \mu_n$ for $n > x_2^*$, there is a restoring force so that $x_2^*$ is a stable equilibrium. Thus, the presence of passive cells impeding T cell elimination in proportion to their abundance generates an equilibrium point $x_2^*$ in direct proportion to their number. Because $x_2^*$ is always stable, cancer escape is impossible, unless the number of passive immunomodulatory cells exceeds the clinically observable tumor size scaled by $(\delta - r)/r$. Phenomenologically, cancer sizes above $x_2$ result in increasing immune encounters that ultimately enhance the death rate above the growth rate.

Moreover,

$$\frac{\partial \xi}{\partial M} = \frac{x^2}{(M+x)^2} > 0, \quad \text{for } x > 0.$$

As such, increases in the inhibitory signal $M$ increases both the number of cancer cells at stable equilibrium and the barrier for the population to become eliminated. We remark that this stability, together with the fact that all population sizes can reach the absorbing extinction state with nonzero probability, implies that the likelihood of ultimate cancer extinction is 1. The expected absorption time to zero, $T_0$, is well-known [31] and given by:

$$\mathbb{E}\left[T_0\right] = \sum_{i=1}^{\infty} \rho_i, \qquad \text{with } \rho_i \equiv \frac{1}{\mu(i)} \prod_{j=1}^{i-1} \frac{\lambda(j)}{\mu(j)}.$$

From equations (S1) and (S2), we have that

$$\rho_i = \frac{\prod_{j=1}^{i-1} rj}{\prod_{j=1}^{i} \delta j \left(\frac{j}{M+j}\right)} = \frac{1}{ri}\left(\frac{r}{\delta}\right)^i \binom{M+i}{i} < \frac{1}{r}\left(\frac{r}{\delta}\right)^i \binom{M+i}{i} \equiv \rho_{max}.$$

We can use the Binomial theorem and the fact that

$$\binom{n}{k} = (-1)^k \binom{k-(n+1)}{k},$$

to show that the expected absorption time is finite. Namely,

$$\mathbb{E}\left[T_0\right] < \sum_{i=0}^{\infty} \rho_{max} = \frac{1}{(1-r/\delta)^{M+1}} < \infty.$$

Without any additional assumption on the value of $M$, these predicted dynamics imply that, for all reasonable parameter choices (i.e. $\delta > r$, $M > 0$), a cancer population subject to passive immunomodulation will never escape to arbitrarily large sizes but will instead reach an equilibrium state. Moreover, given finite time, fluctuations in the population size will ultimately result in disease elimination. For cases with very large $M$, the cancer population is predicted to stabily exist indefinitely. Thus, the theory predicts that passive inhibition alone always leads to population control, and a description of escape therefore requires additional and active immune inhibition. Before considering such models, we first detail the existence and stability of equilibrium states for generalized inhibition functions $f$.

#### S5 Sufficient conditions for equilibria and their stability

For this section, we will represent the immunomodulation function by $f(n)$. We assume that the death rate takes the following general functional form:

$$\mu_n = \delta n f(n); \qquad 0 \leq f(n) \leq 1, \;\; f(0) = f_0 \in \mathbb{R}, \tag{S4}$$

where the immune inhibitory response, $f(x)$, is continuous. The net growth rate becomes

$$\xi = x\big(r - \delta f(x)\big) \tag{S5}$$

Here, equilibrium points are again found by solving $\xi = 0$. Stable points occur whenever $\xi' < 0$, while unstable and semi-stable points occur whenever $\xi' \geq 0$. Clearly, $x^* = 0$ is an equilibrium point, and if $f_0 = 0$ it is always unstable since

$$\xi'(0) = r - \delta f_0. \tag{S6}$$

Additional equilibria are members of the set $E_f = \{x^* > 0 : f(x^*) = r/\delta\}$. For $x^* \in E_f$,

$$\xi'(x^*) = r - \delta f(x^*) - \delta x^* f'(x^*) = -\delta x^* f'(x^*). \tag{S7}$$

The stability of $x^* > 0$ is therefore determined by $f'|_{x^*}$. In particular, each $x^* \in E_f$ for which $f$ is decreasing is unstable . Moreover, ultimate population escape or control depends on the stability of the lead term, $x^*_M = \sup E_f$. If $f'(x^*_M) \leq 0$, then the population tends to escape immune control once its size exceeds $x^*_M$. We remark that in Sec. S4, $x^*_M$ was always stable since $f$ was monotonic increasing.

We are primarily interested in understanding the effects of possibly complicated immune inhibitory responses on the ultimate escape or elimination of a population. The asymptotic behavior of $f$ affects the tendency of $\xi$ and is particularly relevant for a cancer which is not immediately detected by the immune system until larger cell sizes. The behavior may be partitioned into three cases based on the following asymptotic behavior of $f$:

$$f \simeq f_\infty - \beta / x^{1+\varepsilon} \tag{S8}$$

**Case I: $f_\infty > r/\delta$.** Here, $\xi \to -\infty$ and so population control occurs. Note that systems for which $f \to 1$ are subsets of this case, including passive immunomodulation case studied initially (Sec. S4). Immune enhancement, though not explicitly considered here, occurs whenever $f > 1$ and is described by this case.

**Case II: $f_\infty < r/\delta$.** Here, $\xi \to \infty$, leading to population escape. This case includes $f \to 0$, where active immunomodulation (studied in Sec. S7) leads to additional detection impairment for larger cancer population sizes.

**Case III: $f_\infty = r/\delta$.** In this special case the rate of approach to $f_\infty$ matters. By Eqs. S5,S8, if $f \simeq r/\delta - \beta/x^{1+\varepsilon}$, the behavior may be subdivided:

a. If $\varepsilon > 0$ then $\xi$ decreases (resp. increases) to 0 for $\beta > 0$ (resp. $\beta < 0$).

b. If $\varepsilon = 0$ then $\xi \to \beta\delta$.

c. If $-1 < \varepsilon < 0$ then $\xi \to -\infty$ (resp. $\xi \to \infty$) if $\beta > 0$ (resp. $\beta < 0$).

To conclude, the long-term behavior $f_\infty$ relative to $r/\delta$ determines whether a cancer population may escape upon reaching a critical size $x^*_M$. We remark that Case III requires that the long-term tendency in immune impairment is precisely $r/\delta$, which is highly unlikely. Therefore, we proceed motivated by Cases I and II.

### S6 Diffusion Approximation, Absorption Probabilities and Mean Explosion Time

Diffusion approximation is a technique used to approximate the behavior of Markov processes. In our case, the birth death model is a Markov process defined under continuous time and discrete state space. The diffusion approximation provides a continuous state space approximation of birth-death processes' dynamics. Furthermore, such approximation allows us to cast the discrete model under a stochastic differential equation framework, thus making it easier to derive mathematical results to study the behavior of the process. In what follows, we look into analyzing the ultimate extinction probability and the mean explosion time of the birth-death process to extinction via its diffusion approximation. We show the diffusion approximation for the passive immodulation case with the understanding that the mathematical framework can be used for a wider class of birth death processes where both growth and death rates are Lipshitz with respect to the population size.

#### S6.1 Diffusion Approximation

For this subsection, we follow the exposition by Ethier and Kurtz in [32]. Consider a birth-death process, $X(t)$, on the natural numbers with the transition probabilities given by

$$\mathbb{P}\left(X(t+\Delta t) = k+l \,|\, X(t) = k\right) = q_{k,k+l}\Delta t, \quad k, l \in \mathbb{Z}, \tag{S9}$$

where $q_{k,k+l}$ is the transition rate between the state $k$ to the state $k+l$. For our case, the transition rates are population dependent and the death rate is non-linear. Namely,

$$q_{k,k+l} = \begin{cases} \lambda_k = rk, & l = 1, \\ \mu_k = \delta k f(k : M), & l = -1, \\ 0, & \text{otherwise,} \end{cases} \tag{S10}$$

where $r$ and $\delta$ are per-cell birth and death rates respectively. Let $N > 0$ denote the maximal system size of the state space, determined by clinically detectable disease or death. We rewrite the transition rates to be density dependent. In addition, the immodulation function, $f$, changes according to the population size but can easily be written in terms of population density. Namely,

$$q^N_{k,k+l} = \begin{cases} \lambda(k) = Nr\dfrac{k}{N} \triangleq N\beta_1(\dfrac{k}{N}), & l = 1, \\ \mu(k) = N\delta\dfrac{k}{N}f(\dfrac{k}{N} : \dfrac{M}{N}) \triangleq N\beta_{-1}(\dfrac{k}{N}), & l = -1 \\ 0, & \text{otherwise.} \end{cases} \tag{S11}$$

We construct a modified Markov process based on the original Markov process with this system size "limitation". Let $N > 0$, then define $\tilde{X}^N(t)$ as the Markov process with the transition rates given by Eq. (S11). Then, we have the semi-group operator associated with $\tilde{X}^N(t)$ given by

$$A^N g(x) = \begin{cases} \displaystyle\sum_l N\beta_l(\frac{x}{N})\left(g(x+l) - g(x)\right), & x \in \mathbb{Z}, \\ 0, & \text{otherwise,} \end{cases} \tag{S12}$$

where $g$ is a bounded function. Before we proceed on the approximation, we have to introduce the idea of martingales. Martingales are stochastic processes for which the present value is the best approximation for the future. Namely, given a random process $X_t$ with finite expectation and $\mathcal{F}_t$, the filtration or information of the history of a random process $X_s$ upto time $t$, the martingale property is defined as

$$\mathbb{E}\left[X_t \,|\, \mathcal{F}_s\right] = X_s \qquad \forall s \le t$$

Following Ethier and Kurtz [32], $X_t$ and $\mathcal{F}_t$ is a solution to a martingale problem for the semigroup operator $A$ if

$$h(X_t) - H(X_0) - \int_0^t A(h)(X_s)ds$$

where $h$ is a function in the domain of $A$, is a martingale with respect to $\mathcal{F}_t$. This enables us to explicitly write down an analytical expression for the Markov process in terms of another probabilistic object: Poisson processes. Namely, via Theorem 4.1 in Chapter 6 of [32], we have that the stochastic process given by

$$\tilde{X}^N(t) = \tilde{X}^N(0) + \sum_l lY_l \left( N \int_0^t \beta_l \left( \frac{\tilde{X}^N(s)}{N} \right) ds \right), \tag{S13}$$

where $Y_l$ is a standard Poisson process, is a solution for the martingale problem for $A^N$. Furthermore, the above equality holds almost surely for $t$ such that $t \in [0, \tau]$ where

$$\tau = \inf\{t > 0 \,|\, X(t) = \infty\}.$$

We define

$$F(x) = \sum_l l\beta_l(x) \tag{S14}$$

and,

$$X^N(t) = \frac{\tilde{X}^N(t)}{N}. \tag{S15}$$

Note that, we have $F(x) = \beta_1(x) - \beta_{-1}(x)$ for birth-death processes. We use equations S15 and S13 in conjunction to get

$$\begin{aligned} X^N(t) &= X^N(0) + \frac{1}{N}\sum_l lY_l \left( N \int_0^t \beta_l \left( X^N(s) \right) ds \right) \\ &= X^N(0) + \frac{1}{N}\sum_l l \left( Y_l \left( N \int_0^t \beta_l \left( X^N(s) \right) ds \right) - N \int_0^t \beta_l \left( X^N(s) \right) ds \right) \\ &+ \sum_l l \int_0^t \beta_l \left( X^N(s) \right) ds \end{aligned}$$

Consider $\tilde{Y}(u) = Y(u) - u$. As the Poisson process centered around its expectation, $\tilde{Y}(u)$ is a martingale with variation $u$. Hence, we have the following equality,

$$X^N(t) = X^N(0) + \frac{1}{N}\sum_l l\tilde{Y}_l \left( N \int_0^t \beta_l \left( X^N(s) \right) ds \right) + \int_0^t F(X^N(s))ds. \tag{S16}$$

We explore two different methods in obtaining our diffusion approximation. In the first method, we construct the other side of the approximation and then show that the two approximations are the same for large $N$. To that extent, heuristically, we use Eq. (S16) to find the generator for $X^N(t)$. We restrict the function $g$ as in Eq. (S12) to twice differentiable functions, expand $g$ to its Taylor series and then drop the higher degree terms. Doing so, we get the following infinitesimal generator,

$$B^N g(x) = \frac{1}{2N}\sum_{i,j} G_{i,j}(x)\partial_i\partial_j g(x) + \sum_i F_i(x)\partial_i g(x) \tag{S17}$$

where $G(x) = \sum_l ll^T \beta_l(x) = \beta_1(x) + \beta_{-1}(x)$.

Assuming the equation obtained as a solution of the martingale problem for $B^N$ is unique, from Theorem 5.1 of chapter 6 of [32], the solution of the martingale problem can be obtained as a solution of the following equation

$$Z^N(t) = X^N(0) + \frac{1}{N}\sum_l lW_l\left(N\int_0^t \beta_l\left(Z^N(s)\right)ds\right) + \int_0^t F(Z^N(s))ds \tag{S18}$$

where, $W_l$ are independent standard Brownian motions and, $t$ is less than the first infinity of jumps. Both $\beta_l$ and $F(x)$ are Lipschitz and as such, from Theorem 3.1, Chapter 11 of [32], $X^N(t)$ and $Z^N(t)$ are close asymptotically for large $N$. From Theorem 5.3 of Chapter 6 in [32], we have that the above diffusion approximation can be given as a solution to the stochastic differential equation defined by,

$$Z^N(t) = X^N(0) + \frac{1}{\sqrt{N}}\sum_l \int_0^t l\sqrt{\beta_l\left(Z^N(s)\right)}dW_l(s) + \int_0^t F(Z^N(s))ds. \tag{S19}$$

.

As such, we have that the diffusion approximation of $X^N(t)$ is given by a diffusion with mean $F(x)$ and variance $\frac{1}{N}G(x)$:

$$Z^N(t) = Z^N(0) + \frac{1}{\sqrt{N}}\int_0^t \sqrt{G\left(Z^N(s)\right)}dW(s) + \int_0^t F(Z^N(s))ds. \tag{S20}$$

###### S6.1.1 Alternate way of deriving diffusion using local martingales

In the previous subsection, we see that the Poisson processes in Eq. (S16) was approximated by Brownian motion in Eq. (S19). We demonstrate another method to perform such approximation. For this, we start off with Eq. (S16) and look at

$$\frac{1}{N}\sum_l l\tilde{Y}_l\left(N\int_0^t \beta_l\left(X^N(s)\right)ds\right),$$

where, $\tilde{Y}_l$ is a mean zero Poisson process with unit rate. This means that we can rewrite

$$\tilde{Y}_l\left(N\int_0^t \beta_l\left(X^N(s)\right)ds\right) = \int_0^t \sqrt{N\beta_l\left(X^N(s)\right)}\,dM(s)$$

where $M(t)$ is a local martingale with

$$M(t) = \int_0^t \frac{1}{\sqrt{N\beta_l\left(X^N(t)\right)}}\tilde{Y}_l\left(N\int_0^t \beta_l\left(X^N(s)\right)ds\right)$$

Using the fact that $\tilde{Y}$ is a martingale, we can evaluate the quadratic variation of $M(t)$. Namely,

$$\begin{aligned}
[M]_t &= \int_0^t \frac{1}{N\beta_l\left(X^N(r)\right)}d\left[\tilde{Y}_l\left(N\int_0^t \beta_l\left(X^N(s)\right)ds\right)\right] \\
&= \int_0^t \frac{1}{N\beta_l\left(X^N(r)\right)}dY_l\left(N\int_0^t \beta_l\left(X^N(s)\right)ds\right) \\
&\implies t,
\end{aligned} \tag{S21}$$

where we have used the fact that the quadratic variation of a compensated poison process is the original Poisson process itself. Thus, $M(t)$ is a local martingale and has quadratic variation equal to $t$ (in probability). Via Levy's characterization, $M(t)$ is Brownian motion. This gives us

$$\frac{1}{N}\sum_{l} l\tilde{Y}_l\left(N\int_0^t \beta_l\left(X^N(s)\right)ds\right) \implies \frac{1}{\sqrt{N}}\sum_{l} l\int_0^t \sqrt{N\beta_l\left(X^N(s)\right)}\,dW_l(s)$$

where $W_l$ are independent Brownian motions. Using this approximation and a result about weak convergence of probability measures in [33], we arrive at the same stochastic differential equation as in Eq. (S19).

#### S6.2 Absorption Probabilities to Extinction

For this section, we follow the exposition provided by Tan in his book [34]. From the previous section, we are looking for absorption probabilities for a diffusion process with mean $F(x)$ and variance $\frac{1}{N}G(x)$.

Let $u_a(p,t)$ be the probability that the diffusion process $Z^N(t)$ starting at $p$ is absorbed into $a$ before or at time $t$. For formally, we define the following transformation of the conditional pdf of $Z^N$,

$$u_a(p,t) = u(x;s,t) = \int_0^1 f(x,z;s,t)\delta(z-a)dz = f(x,a;s,t)$$

where $f$ is the conditional pdf of the diffusion process $Z^N$. In our case, we are looking for extinction probabilities which is absorption to zero ($a=0$). From Theorem 6.2 of [34], we have that $u_0(p,t)$ satisfies the Kolmogorov backward equation. Namely,

$$\frac{\partial}{\partial t}u_0(p,t) = F(p)\frac{\partial}{\partial p}u_0(p,t) + \frac{1}{2N}G(p)\frac{\partial^2}{\partial p^2}u_0(p,t) \tag{S22}$$

We define $U_0(p) = \lim_{t\to\infty} u_0(p,t)$ as the ultimate probability of elimination. Taking the limit as $t$ grows large on both sides of Eq. (S22), we get

$$F(p)\frac{d}{dp}U_0(p) + \frac{1}{2N}G(p)\frac{d^2}{dp^2}U_0(p) = 0 \tag{S23}$$

with $U_0(0) = 1$ and $U_0(1) = 0$. Using an integrating factor, it can be shown that the the solution of (S23) is given by

$$U_0(p) = 1 - \frac{\int_0^p \phi(x)dx}{\int_0^1 \phi(x)dx}, \tag{S24}$$

where $\phi$ is the scale function given by

$$\phi(x) = \exp\left\{-2N\int_0^x \frac{F(y)}{G(y)}dy\right\}. \tag{S25}$$

For more information, see Chapter 7 of [34]. In finding the extinction probabilities, we use scale function alongside the parameters for the diffusion approximation defined in Eq. (S11). To that extent

$$F(x) = rx - \frac{\delta x^2}{x+\frac{M}{N}}, \qquad G(x) = rx + \frac{\delta x^2}{x+\frac{M}{N}}.$$

We focus on the exponential term in Eq. (S25) :

$$\begin{aligned}
-2N\int_0^x \frac{F(y)}{G(y)}dy &= -2N\int_0^x \frac{ry\left(y+\frac{M}{N}\right)-\delta y^2}{ry\left(y+\frac{M}{N}\right)+\delta y^2}dy,\\
&= -2N\int_0^x \frac{y(r-\delta)+\frac{Mr}{N}}{y(r+\delta)+\frac{Mr}{N}}dy,\\
&= -2N\int_0^x \frac{ay+b}{cy+b}dy,
\end{aligned}$$

where $a \triangleq (r-\delta)$, $b \triangleq \frac{Mr}{N}$, and $c \triangleq r+\delta$ for the ease of computation. Then, we have

$$\begin{aligned}
-2N\int_0^x \frac{F(y)}{G(y)}dy &= -2N\left(\frac{b(c-a)\log(b+cy)+acy}{c^2}\bigg|_{y=0}^{y=x}\right),\\
&= \frac{-4Nb\delta\left(\log(b+cx)-\log(b)\right)-2Nacx}{c^2},\\
&= \frac{-4Nb\delta\log(b+cx)}{c^2}+\frac{4Nb\delta\log(b)}{c^2}-\frac{2Nax}{c},\\
&= \log(b+cx)^{\frac{-4Nb\delta}{c^2}}+\log b^{\frac{4Nb\delta}{c^2}}-\frac{2Nax}{c}.
\end{aligned}$$

Taking the exponential of both sides, we get

$$\phi(x) = (b+cx)^{\frac{-4Nb\delta}{c^2}}\, b^{\frac{4Nb\delta}{c^2}}\, e^{-\frac{2Nax}{c}}.$$

We take the integral with respect of $x$ from 0 to $p$ to get,

$$\begin{aligned}
\int_0^p \phi(x)dx &= \int_0^p (b+cx)^{\frac{-4Nb\delta}{c^2}}\, b^{\frac{4Nb\delta}{c^2}}\, e^{-\frac{2Nax}{c}}dx,\\
&= b^{\frac{4Nb\delta}{c^2}}\int_0^p (b+cx)^{\frac{-4Nb\delta}{c^2}}\, e^{-\frac{2Nax}{c}}dx.
\end{aligned}$$

To further aid in computation, let us redefine some new constants:

$$\alpha = \frac{4Nb\delta}{c^2} = \frac{4Mr\delta}{c^2} \quad \text{and } \eta = \frac{2Na}{c}.$$

Furthermore, let $u = \frac{\eta}{c}(b+cx)$. Then, via $u$-substitution, we have

$$\begin{aligned}
\int_0^p \phi(x)dx &= b^\alpha \int_0^p (b+cx)^{-\alpha}e^{-\eta x}dx\\
&= b^\alpha \int_0^p \left(\frac{cu}{\eta}\right)^{-\alpha} e^{-u+\frac{b\eta}{c}}du.\frac{1}{\eta}\\
&= \left(\frac{\eta b}{c}\right)^\alpha \frac{1}{\eta}e^{\frac{b\eta}{c}}\int_0^p u^{-\alpha}e^{-u}du\\
&= \left(\frac{\eta b}{c}\right)^\alpha \frac{1}{\eta}e^{\frac{b\eta}{c}}\int_{\frac{\eta b}{c}}^{\frac{\eta b}{c}+\eta p} u^{(1-\alpha)-1}e^{-u}du\\
&= \left(\frac{\eta b}{c}\right)^\alpha \frac{1}{\eta}e^{\frac{b\eta}{c}}\left[\Gamma(1-\alpha,\frac{\eta b}{c})-\Gamma(1-\alpha,\frac{\eta b}{c}+\eta p)\right],
\end{aligned}$$

where $\Gamma$ is the incomplete upper gamma function given by

$$\Gamma(s, x) = \int_x^\infty t^{s-1} e^{-t} dt. \tag{S26}$$

Finally, we can obtain the ultimate extinction probability in Eq. (S24):

$$\begin{aligned} U_0(p) &= 1 - \frac{\int_0^p \phi(x)dx}{\int_0^1 \phi(x)dx} \\ &= 1 - \frac{\Gamma(1-\alpha, \frac{\eta b}{c}) - \Gamma(1-\alpha, \frac{\eta b}{c} + \eta p)}{\Gamma(1-\alpha, \frac{\eta b}{c}) - \Gamma(1-\alpha, \eta(\frac{b}{c}+1))} \\ &= \frac{\Gamma(1-\alpha, \frac{\eta b}{c} + \eta p) - \Gamma(1-\alpha, \eta(\frac{b}{c}+1)}{\Gamma(1-\alpha, \frac{\eta b}{c}) - \Gamma(1-\alpha, \eta(\frac{b}{c}+1))}. \end{aligned}$$

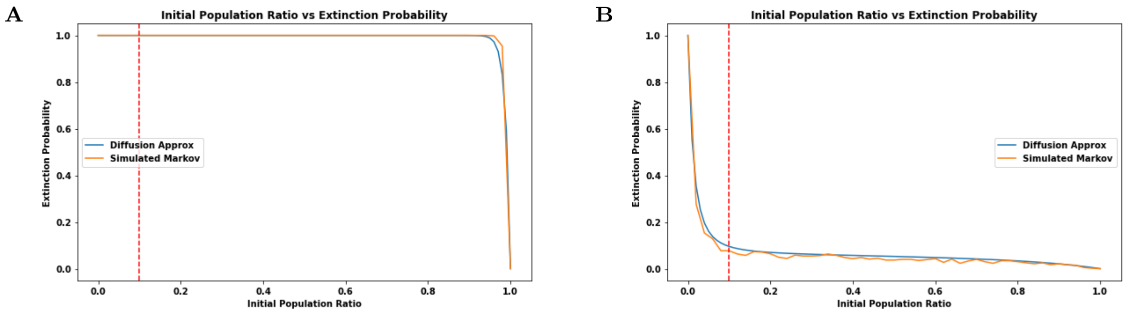


Figure S1: Ultimate Extinction Probability as a function of initial population ratio. We compare the dynamics of the simulated birth-death process and its diffusion approximation. **A.** $N = 100,\ r = 0.005,\ d = 1$ **B.** $N = 50,\ r = 0.45,\ d = .55$.

Here, the ultimate escape probability when starting at the population ratio $p$ can be found to be

$$\mathbb{P}_{esc}(p) = 1 - U_0(p) \tag{S27}$$

In Fig. S1, we applied Gillespie algorithm to simulate the underlying Markov chain and compared the elimination probability with the diffusion approximation given in (S24). In both cases, we find that unless the initial population ratio is close to unity, the process will be predicted to be eliminated. This is in line with what we expected for the passive immunization knowing that the process was predicted to die out in the discrete space state Markov chain for large $N$ when $r \ll \delta$.

#### S6.3 Mean Escape Time

Given the stochastic differential equation obtained in the diffusion approximation from Eq. (S20), we look into another interesting property: average escape time. To that extent, define the stopping time,

$$\tau = \inf \{t \geq 0 \,|\, X(t) \geq 1\}. \tag{S28}$$

Let $g$ be a differentiable function that is to be determined later and consider $g\left(X_t\right)$. By Itô's Lemma, we have

$$\begin{aligned} dg(X_t) &= g'(X_t)dX_t + \frac{1}{2}g''(X_t)\left(dX_t\right)^2 \\ &= \left[F(X_t)g' + \frac{1}{2N}g''G(X_t)\right]dt + \frac{1}{\sqrt{N}}g'(X_t)\sqrt{G(X_t)}dB_t \end{aligned}$$

where $g'$ amd $g''$ represent differential of the function $g$ with respect to the $x$ variable evaluated at $X_t$. We integrate the SDE from 0 to $\tau$ to get

$$g(X_\tau) - g(X_0) = \int_0^\tau \left[F(X_s)g'(X_s) + \frac{1}{2N}g''(X_s)G(X_s)\right]ds + \int_0^\tau \frac{1}{\sqrt{N}}g'(X_s)\sqrt{G(X_s)}dB_s$$

We take the expectation of both sides and apply the Optional Stopping theorem to get

$$g(1) - g(X_0) = \mathbb{E}\left[\int_0^\tau \left(F(X_s)g'(X_s) + \frac{1}{2N}g''(X_s)G(X_s)\right)ds\right]$$

Suppose that $g$ satisfies the following ordinary differential equation with boundary values:

$$\begin{cases} F(X_t)g' + \frac{1}{2N}g''G(X_t) = -1, \\ g(1) = 0, \quad g'(1) = 0. \end{cases} \tag{S29}$$

Then we can use this function and the initial population ratio to obtain the average time for explosion. Namely,

$$\mathbb{E}\left[\tau \mid X_0\right] = g(X_0).$$

Thus, we pivot to solving the differential equation presented above. We can rewrite Eq. (S29) in the following way:

$$\begin{cases} g'' + p(x)g' = q(x), \\ g(1) = 0, \quad g'(1) = 0 \end{cases} \tag{S30}$$

where,

$$p(x) = \frac{2N\left(\left(r-\delta\right)x + \frac{rM}{N}\right)}{\left(r+\delta\right)x + \frac{Mr}{N}}, \quad q(x) = \frac{-2N(x + \frac{M}{N})}{\left(r+\delta\right)x^2 + \frac{Mrx}{N}}.$$

Note that both $p$ and $q$ are continuous on the interval $(0,1)$ and as such, there exists a unique solution to the second order non-homogeneous linear differential equation given by Eq. (S30). We start from the homogeneous form to obtain the general solution.

$$\begin{aligned} g'' + p(x)g' &= 0 \\ e^{-\int_x^1 p(v)dv}(g'' + p(x)g') &= 0 \\ e^{-\int_x^1 p(v)dv}g'' + p(x)e^{-\int_x^1 p(v)dv}g' &= 0 \\ (e^{-\int_x^1 p(v)dv}g')' &= 0 \\ g' &= -c_2 e^{\int_x^1 p(v)dv} \\ g(x) &= c_2\int_x^1 e^{\int_y^1 p(v)dv}dy + c_1 \end{aligned}$$

where we have integrated from some arbitrary initial value $x \in (0,1)$ to the final value $X(\tau) = 1$. Thus, we have the two homogeneous solutions given by

$$g_1(x) = 1, \qquad g_2(x) = \int_x^1 e^{\int_y^1 p(v)dv} dy.$$

We calculate the Wronskian for these two solutions:

$$\begin{aligned} W(x) &= g_1 g_2' - g_1' g_2 \\ &= g_2' \\ &= -e^{\int_x^1 p(v)dv} \\ &\neq 0 \, \forall \, x \in [0,1] \end{aligned}$$

This means that the two solutions are fundamental solutions for the general case. We now use these general case solutions and method of variation of parameters to obtain the particular solution for the non-homogeneous case. It is well known that the particular solution will have the form

$$\begin{aligned} g_p(x) &= -g_1 \int_x^1 \frac{g_2(v)q(v)}{W(v)} dv + g_2 \int_x^1 \frac{g_1(v)q(v)}{W(v)} dv \\ &= -\int_x^1 \frac{g_2(v)q(v)}{W(v)} dv + g_2(x) \int_x^1 \frac{q(v)}{W(v)} dv \end{aligned} \tag{S31}$$

For the first term in the right hand side of the above equation,

$$\begin{aligned} -\int_x^1 \frac{g_2(v)q(v)}{W(v)} dv &= -\int_x^1 \frac{\int_v^1 e^{\int_y^1 p(z)dz} dy}{-e^{\int_v^1 p(z)dz}} \, q(v)dv \\ &= \int_x^1 \int_v^1 \frac{e^{\int_y^1 p(z)dz}}{e^{\int_v^1 p(z)dz}} dy \; q(v)dv \\ &= \int_x^1 \int_v^1 e^{-\int_v^y p(z)dz} dy \; q(v)dv \end{aligned} \tag{S32}$$

Similarly from (S31),

$$\begin{aligned} \int_x^1 \frac{g_2(x)q(v)}{W(v)} dv &= \int_x^1 \frac{\int_x^1 e^{\int_y^1 p(z)dz}}{-e^{\int_v^1 p(z)dz}} dy \; q(v)dv \\ &= -\int_x^1 \int_x^1 e^{\int_y^1 p(z)dz - \int_v^1 p(z)dz} dy \; q(v)dv \\ &= -\int_x^1 \int_x^1 e^{-\int_v^y p(z)dz} dy \; q(v)dv \end{aligned} \tag{S33}$$

Combining Eqs. (S31),(S32) and, (S33), we get

$$\begin{aligned} g_p(x) &= \int_x^1 \int_v^1 e^{-\int_v^y p(z)dz} dy \; q(v)dv - \int_x^1 \int_x^1 e^{-\int_v^y p(z)dz} dy \; q(v)dv \\ &= \int_x^1 \left[ \int_v^1 e^{-\int_v^y p(z)dz} dy - \int_x^1 e^{-\int_v^y p(z)dz} dy \right] q(v)dv \\ &= -\int_x^1 \int_x^v e^{-\int_v^y p(z)dz} dy \; q(v)dv. \end{aligned}$$

Thus, the general form for our solution is given by

$$g(x) = c_1 g_1 + c_2 g_2 + g_p \tag{S34}$$

$$= c_1 + c_2 \int_x^1 e^{\int_y^1 p(v)dv} dy - \int_x^1 \int_x^v e^{-\int_v^y p(z)dz} dy \, q(v) dv, \tag{S35}$$

where $c_1$ and $c_2$ depend on the boundary conditions for $g(1)$ and $g'(1)$. From Eq. (S35) we can see that, $g(1) = 0$ implies that $c_1 = 0$. Furthermore, using FTC and Leibniz Integral Rule we get

$$g'(x) = -c_2 e^{\int_x^1 p(v)dv} + \int_x^1 e^{-\int_v^x p(z)dz} q(v) dv + \int_x^1 e^{-\int_x^y p(z)dz} q(x) dy.$$

As such, $g'(1) = 0$ implies that $c_2 = 0$.

Thus, conditional on the fact that $X_0 = x_0 \in (0, 1]$, we can then find the mean time for escape to be given by

$$\mathbb{E}\left[\tau \mid X_0 = x_0\right] = -\int_{x_0}^1 \int_{x_0}^v e^{-\int_v^y p(z)dz} dy \, q(v) dv \tag{S36}$$

Finally, in Fig. 2 of the main text, we plot the escape probability and mean time to escape to compare the results obtained analytically from the diffusion approximation via Eqns. (S27) and (S36) to large scale stochastic simulations of the underlying markov model.

#### S7 Active immunomodulatory participation

Sec. S4 demonstrated that passive immunomodulation exclusively due to the presence of a fixed inhibition signal results in tumor control. In the language of Sec. S5, passive immunomodulation yields $f_\infty = 1 > r/\delta$, resulting in stable population cancer control. While this may reasonably model environments where active immunosuppression is not important, it does not account for observations where the tumor microenvironment itself is actively immunosuppressive. For example, previous studies of the tumor microenvironment suggest that T cell function is further impaired via metabolic dysfunction [35], and that tumor cell glucose-depleted and lactate-rich environment together lead to enhanced Treg suppressor activity [36]. In the context of a dynamic population, these findings provide a tangible mechanism by which inhibitory cells may become more active as the tumor population increases, thus driving an immune permissive microenvironment. In general, active immunomodulatory behavior may be added by altering the immunomodulation function to include additional T cell killing impairment due to increases in the inhibitory signal $M$ or total tumor cells $n$. In contrast with passive environmental barriers to killing, this framework more appropriately characterizes the presence of a fixed amount of inhibiting signal, such as myeloid-derived stem cells or depletion of an exogeneous required metabolite. While there are many such functional dependencies that could describe this, we will for foundational understanding consider two simple cases which build in two scaled contributions to inhibition, $\alpha M$ and $\beta n$, via:

$$f_p(n, M) = \left(\frac{n}{M+n}\right)\left(\frac{1}{\alpha M \cdot \beta n}\right); \qquad f_l(n; M) = \left(\frac{n}{M+n}\right)\left(\frac{1}{\alpha M + \beta n}\right) \tag{S37}$$

We will refer to systems adopting $f_p$ as *product inhibition* and those adopting $f_l$ as *linear inhibition.*

##### S7.1 Product inhibition

$f_p$ yields the net growth rate

$$\xi(x) = rx - \frac{\delta x}{\alpha\beta M\,(x+M)}.$$

with the roots located at

$$x = 0,\ \frac{\delta}{\alpha\beta M r} - M.$$

Then,

$$\xi'(x) = r - \frac{\delta}{\alpha\beta}\left(\frac{1}{x+M}\right)^2.$$

This allows us to analyse at the behavior at the two roots. Hence we have

$$\xi'(0) = r - \frac{\delta}{\alpha\beta M^2}, \qquad \xi'\left(\frac{\delta}{\alpha\beta M r} - M\right) = r\left(1 - \frac{M^2\alpha\beta r}{\delta}\right).$$

We look into two different cases:

- **Case 1:** $\boldsymbol{\frac{\delta}{\alpha\beta r} < M^2}$. Then the rightmost equilibrium point is at $x = 0$.

$$\begin{aligned}\xi'(0) &= r - \frac{\delta}{\alpha\beta M^2},\\ &= r - \frac{r}{M^2}\frac{\delta}{\alpha\beta r},\\ &> r - \frac{r}{M^2}M^2,\\ &= 0\end{aligned}$$

- **Case 2: $\boldsymbol{\frac{\delta}{\alpha\beta r} > M^2}$**. Then we have two positive roots. Then,

$$\begin{aligned}\xi'\left(\frac{\delta}{\alpha\beta M r} - M\right) &= r\left(1 - \frac{M^2\alpha\beta r}{\delta}\right),\\ &> r\left(1 - \frac{M^2}{M^2}\right),\\ &= 0.\end{aligned}$$

While,

$$\begin{aligned}\xi'(0) &= r - \frac{\delta}{\alpha\beta M^2},\\ &= r - \frac{r}{M^2}\frac{\delta}{\alpha\beta r},\\ &< r\left(1 - \frac{M^2}{M^2}\right),\\ &= 0.\end{aligned}$$

Thus, in the first case, the singular positive root is an unstable equilibrium. In the latter case, the root at 0 is absorbing while the positive root is an unstable equilibrium. We illustrate that fact with figures below.

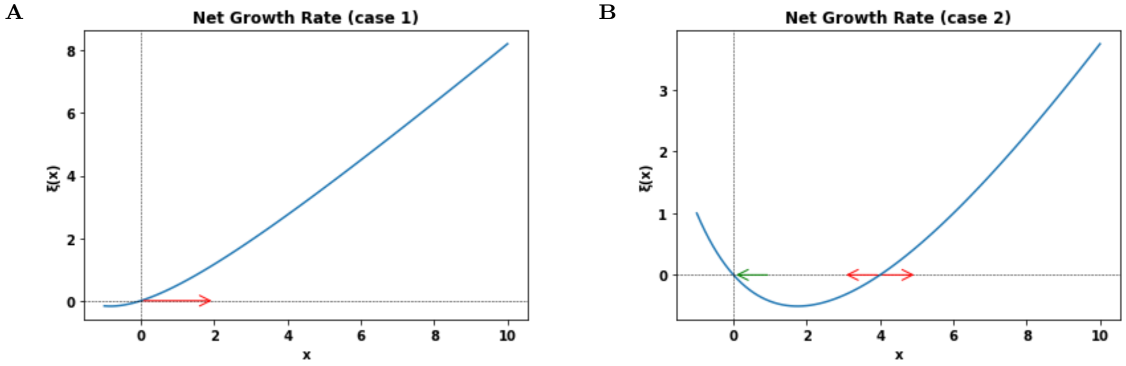


Figure S2: Schematic net Growth Dynamics for Product Inhibition depending on the parametrization of $\alpha, \beta, r, \delta$, and $M$. **A.** When $\delta < \alpha\beta r M^2$ then under product inhibition, we obtain one unstable state at origin **B.** Conversely, when $\delta > \alpha\beta r M^2$, then we have one stable state at origin and a unstable strictly positive root. Both cases promote escape and collectively are not equipped for a stable nonzero equilibrium state.

Now we look at the net growth rate with respect to the immodulatory constant. Namely,

$$\frac{\partial\xi(x)}{\partial M} = \frac{\delta x\,(2xM + 1)}{\alpha\beta M^2\,(Mx + 1)^2} > 0, \qquad \text{when } x \geq 0.$$

#### S7.2 Linear inhibition

$f_l$ yields the size-inhomogeneous net-growth rate:

$$\xi(x) = rx - \frac{\delta x^2}{(M + x)(\alpha M + \beta x)}. \tag{S38}$$

As in Section S4, equilibrium points correspond to roots of $\xi$. In addition to $x = 0$, the roots of Eq. S38 are found by solving

$$g(x, \alpha) \equiv \beta x^2 + [(\alpha + \beta) M - \gamma] x + \alpha M^2 = 0, \qquad \text{for } \gamma \equiv \delta/r > 1. \tag{S39}$$

Solutions to $g(x, \alpha)$ depend on its discriminant, $D(\alpha, \beta)$, which can be written as

$$D(\alpha, \beta) = \left[\eta^2 - 2(\alpha + \beta)\eta + (\alpha - \beta)^2\right] M^2, \qquad \text{for } \eta \equiv \gamma/M.$$

$\alpha$ and $\beta$ values relate to the roots, $\mathcal{R}_\xi(\alpha, \beta)$, of $\xi$. Critical values occur when the $D(\alpha, \beta) = 0$ and are determined by the solution to

$$\eta^2 - 2(\alpha + \beta)\eta + (\alpha - \beta)^2 = 0.$$

This occurs for

$$\begin{aligned} \eta &= (\alpha + \beta) \pm \sqrt{(\alpha + \beta)^2 - (\alpha - \beta)^2} \\ &= (\alpha + \beta) \pm 2\sqrt{\alpha\beta}, \end{aligned}$$

so that

$$\sqrt{\eta_-} = \sqrt{\alpha} - \sqrt{\beta}; \qquad \sqrt{\eta_+} = \sqrt{\alpha} + \sqrt{\beta}.$$

To simplify the analysis in the results to follow and without loss of generality, we normalize by $\beta$ so that we consider

$$f_l(n) = \left(\frac{n}{M + n}\right)\left(\frac{1}{\alpha M + n}\right)$$

where now $\alpha$ has been normalized by dividing by $\beta$, $\gamma \equiv \delta/r\beta$, and $\eta \equiv \gamma/M$. In this case, the critical $\alpha$ values yielding a zero discriminant for $\xi$ become

$$\alpha_\pm = \frac{1}{2}\left[2(\eta + 1) \pm \sqrt{4(\eta + 1)^2 - 4(\eta - 1)^2}\right] = (\eta + 1) \pm 2\sqrt{\eta},$$

yielding

$$\alpha_- = \left(\sqrt{\eta} - 1\right)^2; \qquad \alpha_+ = \left(\sqrt{\eta} + 1\right)^2.$$

From this, we consider all possible cases where $\alpha \geq 0$ as follows:

- **Case I: $\boldsymbol{\alpha = 0}$**. Here, $D(0) = (M - \gamma)^2$, so that the roots

$$\begin{aligned} \mathcal{R}_\xi(\alpha = 0) &= \frac{1}{2}\left(\gamma - M \pm \sqrt{(M + \gamma)^2}\right) \\ &= \{0, \gamma - M\}. \end{aligned}$$

  Thus, we have the following sub-cases:

  a) $\boldsymbol{M < \gamma}$. Equilibrium points are $0 = x_1 < x_2 = \gamma - M$, with Eq. S39 giving $g(x, 0) < 0$ for $x \in (x_1, x_2)$, and $g(x, 0) > 0$ for $x > x_2$ so that $x_1 = 0$ is stable and $x_2 = \gamma - M$ is unstable.

  b) $\boldsymbol{M \geq \gamma}$. In this case, the only nonnegative equilibrium point is $x_1 = 0$ and it is unstable since $g(x, 0) = x(x + M - \gamma)$ is positive for $x > 0$.

- **Case II: $\boldsymbol{0 < \alpha < \alpha_-}$:**. In addition to $x = 0$, $D(\alpha) > 0$ gives two additional real-valued roots. Writing the solutions to $g$ and re-arranging $D(\alpha)$ gives

$$x = \frac{M}{2}\left(-b \pm \sqrt{b^2 - 4\alpha}\right); \qquad b \equiv \alpha + 1 - \eta. \tag{S40}$$

  From here, it is clear that $|b| > |(D(\alpha)|$, which leads to the following sub-cases:

a) $\boldsymbol{M < \gamma}$. Note that $\alpha$ is bounded above by $\alpha_-$, so that $b < 2\left(1 - \sqrt{\eta}\right)$. Thus, $\eta > 1$ (equivalently $M < \gamma$) implies $b < 0$ and so $g$ has two positive roots, $x_1$ and $x_2$, for $0 < \alpha < \alpha_-$. The ordered list of roots of $\xi$, is given by $\mathcal{R}_\xi = \{0, x_1, x_2\}$. Since $f$ in this section corresponds to Case IIIb of Sec. S5, the lead root is unstable ($\xi > 0$ for $x > x_2$). Since $\xi$ is continuous and non-constant, it follows that $\xi > 0$ on $(0, x_1)$, $\xi < 0$ on $(x_1, x_2)$. Thus, 0 is an unstable equilibrium point, $x_1$ is stable, and $x_2$ unstable.

b) $\boldsymbol{M \geq \gamma}$. It must be that $b > 0$ in this case, which implies that g has two negative roots. Otherwise, $b \leq 0$ implies that $\alpha \leq \eta - 1 \leq 0$ for $\eta \leq 1$. But this is impossible since $0 < \alpha < \alpha_-$. Therefore, 0 is the only relevant root, and in this case $g \geq x^2 + \alpha M x + \alpha M^2 > 0$ for $x > 0$, thus it is unstable. We note additionally that $\gamma = M$ implies that $\alpha_- = 0$ so that $\alpha < \alpha_-$ is impossible.

- **Case III:** $\boldsymbol{\alpha = \alpha_-}$. Here, $D(\alpha_-) = 0$. Therefore, the double root of $g$, $x_1$, is given by

$$x_1 = \frac{\gamma - (\alpha + 1) M}{2} = \frac{\gamma - \left(\eta - 2\sqrt{\eta} + 2\right) M}{2} = \sqrt{\gamma M} - M.$$

Thus, $\mathcal{R}_\xi = \{0, \sqrt{\gamma M} - M\}$ and we have the following sub-cases:

a) $\boldsymbol{M < \gamma}$. Then $x_1 = \sqrt{\gamma M} - M$ is a positive root. Moreover, since this root corresponds to the minima of $g$ convex , it is a semi-stable point.

b) $\boldsymbol{M \geq \gamma}$. Then the only nonnegative root is 0, which is unstable.

- **Case IV:** $\boldsymbol{\alpha_- < \alpha < \alpha_+}$. Here, $D(\alpha) < 0$, so that the only real root is $x_0 = 0$.

- **Case V:** $\boldsymbol{\alpha = \alpha_+}$. $D(\alpha_-) = 0$. For $g$ with $\alpha = \alpha_+$ and by similar reasoning to Case III, the double root of $g$, $x_1$, is given by

$$x_1 = -\left(\sqrt{\gamma M} + M\right).$$

So that the only relevant root is $x_0 = 0$ unstable.

- **Case VI:** $\boldsymbol{\alpha > \alpha_+}$. Here, $D(\alpha) > 0$, giving two real roots. Note that Eq. S40 holds in this case. Moreover, $\alpha > \alpha_+$ implies that $b > 2(\sqrt{\eta} + 1) > 0$. Thus, the only relevant root is $x_0 = 0$ and unstable.

To summarize: if the number of regulatory cells exceeds the ratio of the killing rate to the birth rate (i.e. $\gamma \leq M$), then $x_0 = 0$ is the only non-negative equilibrium point and it is unstable, resulting in population growth to large sizes. This same scenario also occurs if, despite $\gamma > M$, we have that the active immune modulatory parameter $\alpha$ is in excess of an upper threshold $\alpha_-$ that depends on both $\gamma$ and $M$. If however $\alpha < \alpha_-$ then there exist additional equilibrium states besides $x_0 = 0$. If $\alpha = 0$ gives the only case where $x_0 = 0$ is stable, and there is an unstable, positive equilibrium state. If $\alpha < \alpha_-$, then $x_0 = 0$ becomes unstable, with two additional equilibria, $0 < x_1 < x_2$, with $x_1$ stable and $x_2$ unstable. $\alpha = \alpha_-$ is a critical case with $x_0 = 0$ unstable and $x_1$ semi-stable. These findings are graphically depicted in Fig. S3.

Realistically, a given cancer population and immune microenvironment is expected to feature a fixed per-cell growth rate and level of immune suppression $\alpha$. The interesting behavior occurs whenever there exist stable and unstable states separated by an activation barrier below which the population is kept in check by activated T cells, and above which the microenvironment may repress T cell killing at levels sufficient for cancer escape. This permits the existence of an indolent phenotype, perhaps for very long times, prior to escape or elimination (Fig. S4). Additionally, the likelihood and timing of cancer escape or elimination, as well as the expected population sizes in the stable state, are all determined by $\xi$ in Eq. S38.

**Expected cycling population size:** These sizes are given by the stable nonzero equilibrium point, which exists whenever $\alpha < \left(\sqrt{\delta / r\beta M} - 1\right)^2$, and given by the smaller root of Eq. S40. Since from Eq. S38

$$\frac{\partial \xi}{\partial M} = x^2 \frac{(\alpha + 1)M + (\beta + 1)x}{(M + x^2)(\alpha M + \beta x)^2} > 0, \quad \text{for } x > 0,$$

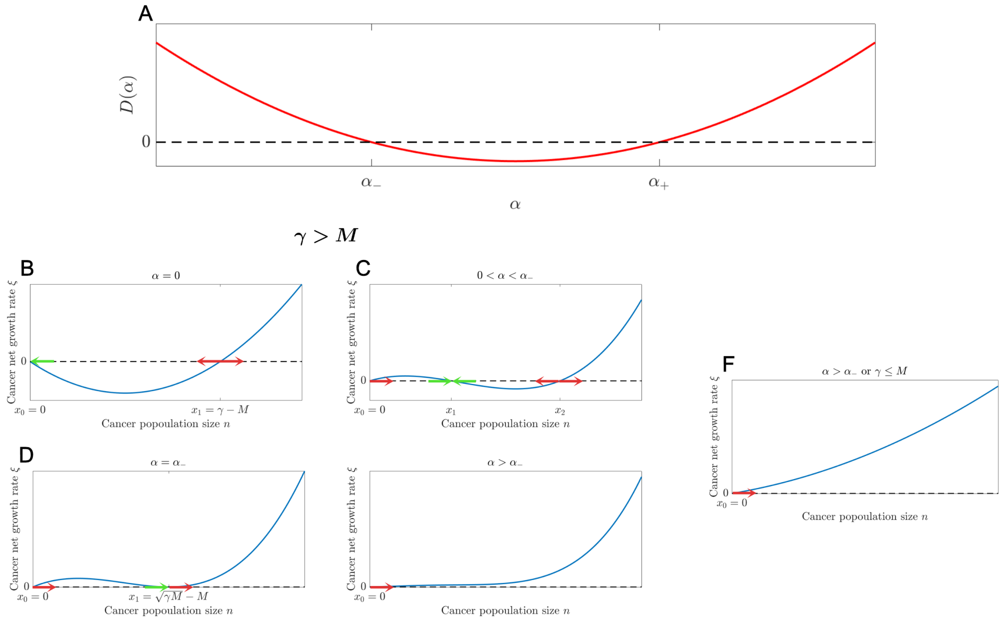

Figure S3: Graphical depiction of the dynamics and equilibrium generated for various levels of immune suppression and regulatory cell sizes. Arrows point in the direction of population tendency (red arrows indicate velocities away from equilibrium points, and green arrows represent velocities toward equilibrium points).

so that the function $\xi(x)$ increases for increasing inhibition. This means that increasing the number of regulatory cells results in three observable behaviors: 1) an increase in the number of cancer cells at stable equilibrium, 2) increases in the lower barrier that determines cancer elimination rates, and 3) decreases in the upper barrier that determines cancer escape rates (Fig. S4A).

**Extinction probability:** By [31], we have that the extinction probability is not guaranteed whenever

$$\sum_{i=1}^{\infty}\prod_{j=1}^{i}\frac{\mu_j}{\lambda_j} < \infty.$$

In this case, we have that

$$\begin{aligned}\sum_{i=1}^{\infty}\prod_{j=1}^{i}\frac{\mu_j}{\lambda_j} &= \sum_{i=1}^{\infty}\prod_{j=1}^{i}\frac{\delta j^2}{rj(M+j)(\alpha M+\beta j)}\\ &\leq \sum_{i=1}^{\infty}\prod_{j=1}^{i}\frac{\delta}{r\beta j} = \sum_{i=1}^{\infty}\left(\delta/r\beta\right)^i/i! = e^{\delta/r\beta}-1 < \infty.\end{aligned}$$

And thus,

$$P_{\text{ext}} = \frac{\sum_{i=N_0}^{\infty}\left(\prod_{j=1}^{i}\frac{\delta j^2}{rj(M+j)(\alpha M+\beta j)}\right)}{1+\sum_{i=1}^{\infty}\left(\prod_{j=1}^{i}\frac{\delta j^2}{rj(M+j)(\alpha M+\beta j)}\right)} \tag{S41}$$

These results are plotted in Fig. S5.

As seen above, the active immune inhibition assumption is an improvement over its passive inhibition counterpart, since this framework permits the existence of stable equilibria which may persist for large times

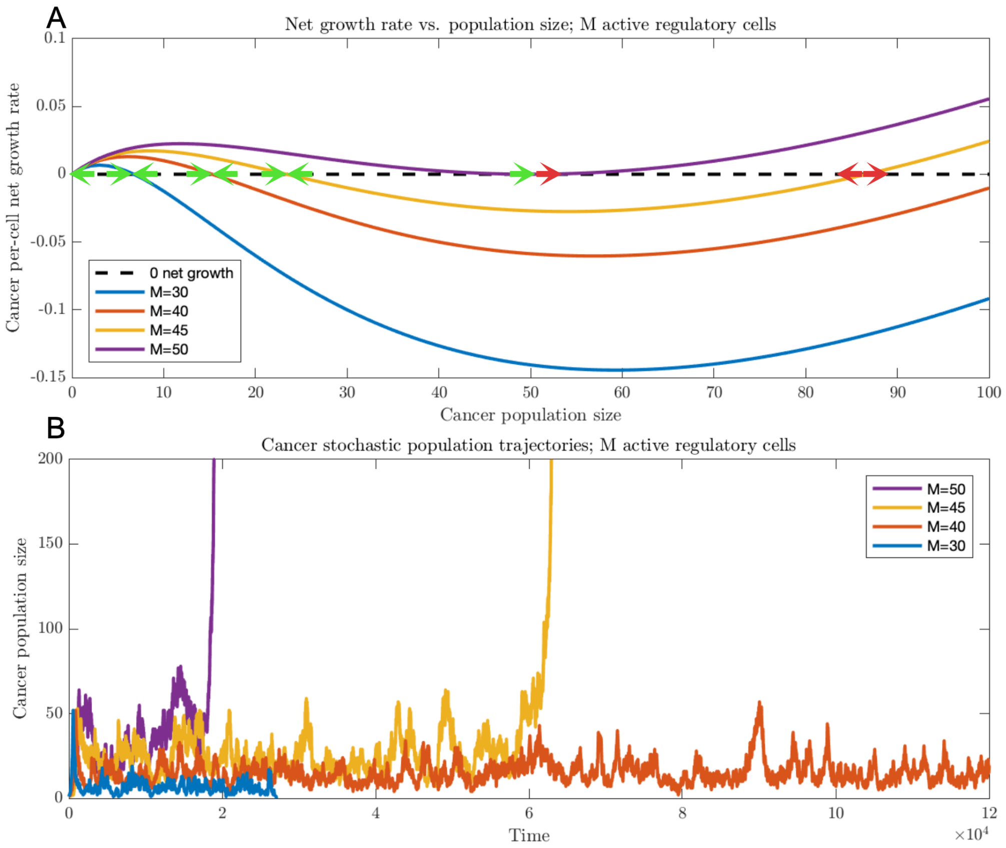


Figure S4: Dynamics of active regulatory cell suppression. A. The cancer population is assumed to follow the dynamics induced by the net growth rate $\xi$ given by Eq. S38 with a variable number of inhibitory cells, $M$ (red arrows indicate velocities away from equilibrium points, and green arrows represent velocities toward equilibrium points); B. Representative stochastic trajectories are depicted (corresponding net growth rates and trajectories sharing the same color; in each case, $r = 0.005$, $d = 1$, giving $\min_M \alpha_-(M) = 1$, with $\alpha = 1$.

until either ultimate escape or elimination occurs. One final task remains, which is to incorporate a dynamic feature into the immunomodulation parameter, which may now itself depend on the cancer population size.

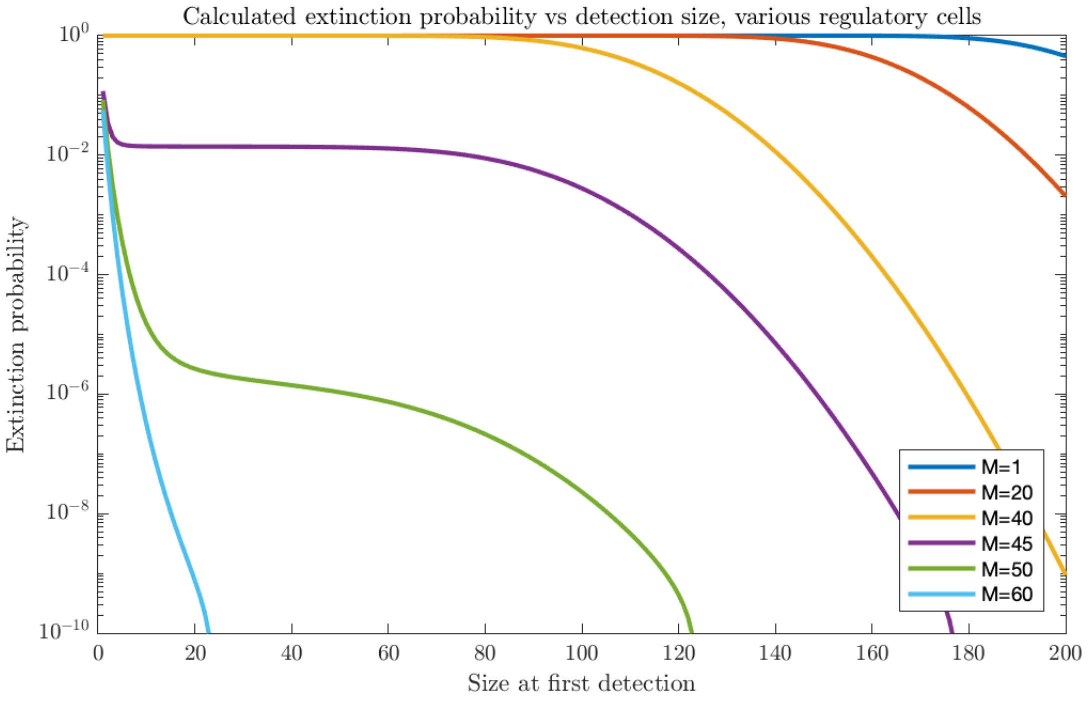


Figure S5: Extinction probability as a function of size at first detection. Extinction probabilities are plotted as a function of initial detection size assuming various numbers of regulatory cells. In all cases, $r = 0.005$, $d = 1$, $\beta = 1$, giving $\min_M \alpha_-(M) = 1$, with $\alpha = 1$ (infinite sums of Eq. S41 were iterated until convergence was achieved with a tolerance of $\varepsilon < 10^{-10}$).

### S8 Dynamic immunomodulatory inhibition

In section S7, we assumed a fixed number of immunomodulatory cells, $M$. Motivated by the observations in Sec. S2.1, we assume that the immunomodulatory parameter may vary during cancer progression as a function of the random cancer population size.

#### S8.1 Conditional hitting probabilities for inhibitory signal size

Toward this end, it will be useful to describe the probability that the cancer population reach a particular size $n$ starting from an initial size $m$ conditional on not first hitting size $d$, denoted by event ${}^{d}E_{n,m}$, for $d < m < n$. Similarly, we denote the event of moving from state $m$ to $n$ by $F_{n,m}$. Then,

$$\begin{aligned}\mathbb{P}\left({}^{d}E_{n+1,n}\right) &= \mathbb{P}\left({}^{d}E_{n+1,n} \mid F_{n+1,n}\right)\,\mathbb{P}\left(F_{n+1,n}\right) + \mathbb{P}\left({}^{d}E_{n+1,n} \mid F^{c}_{n+1,n}\right)\,\mathbb{P}\left(F^{c}_{n+1,n}\right)\\ &= 1\cdot\left(\frac{\lambda_n}{\lambda_n+\mu_n}\right) + \mathbb{P}\left({}^{d}E_{n+1,n-1}\right)\left(\frac{\mu_n}{\lambda_n+\mu_n}\right)\\ &= \nu_n + \eta_{n-1}\,\mathbb{P}\left({}^{d}E_{n,n-1}\right)\,\mathbb{P}\left({}^{d}E_{n+1,n}\right),\end{aligned}$$

where we have defined

$$\nu_n \equiv \frac{\lambda_n}{\lambda_n+\mu_n}; \qquad \eta_n \equiv \frac{\mu_{n+1}}{\mu_{n+1}+\lambda_{n+1}}. \tag{S42}$$

This yields the recursive relation

$$\mathbb{P}\left({}^dE_{n+1,n}\right) = \frac{\nu_n}{1-\eta_{n-1}\,\mathbb{P}\left({}^dE_{n,n-1}\right)}, \quad \text{with } \mathbb{P}\left({}^dE_{d+2,d+1}\right) = \nu_{d+1}. \tag{S43}$$

Put $\gamma_n \equiv \eta_n\nu_n$. Assuming

$$\mathbb{P}\left({}^dE_{n+1,n}\right) = \frac{D_{n-1}}{D_n}\nu_n, \qquad n \geq d+1,$$

then

$$\mathbb{P}\left({}^dE_{n+2,n+1}\right) = \frac{\nu_{n+1}}{1-\eta_n\,\mathbb{P}\left({}^dE_{n+1,n}\right)} = \frac{D_n\nu_{n+1}}{D_n-\gamma_n D_{n-1}}$$

from which it follows that Eq. S43 is satisfied whenever the following relation holds:

$$D_{n+1} = D_n - \gamma_n D_{n-1}, \quad n \geq d+2; \quad \text{with } D_d = D_{d+1} = 1, \tag{S44}$$

We begin with the case where $d = 0$. Note that $D_0$ and $D_1$ are determined by $\mathbb{P}\left({}^0E_{2,1}\right) = \nu_1$ and $\mathbb{P}\left({}^0E_{3,2}\right) = \nu_2/(1-\gamma_1)$.

**Proposition S8.1.** *The explicit solution for $D_n$ defined in Eq.* (S44) *is given by*

$$D_n = 1 + \sum_{k=1}^{\lfloor n/2\rfloor}(-1)^k \sum_{i_1=1}^{n-2k-1}\gamma_{i_1}\left(\sum_{i_2=2+i_1}^{n-2k+1}\gamma_{i_2}\left(\cdots\left(\sum_{i_k=2+i_{k-1}}^{n-1}\gamma_{i_p}\right)\cdots\right)\right) \tag{S45}$$

$$= 1 + \sum_{k=1}^{\lfloor n/2\rfloor}(-1)^k \sum_{\boldsymbol{i}\in\mathcal{S}_k^{(n)}}\prod_{j=1}^{k}\gamma_{i_j}, \tag{S46}$$

*where*

$$\mathcal{S}_k^{(n)} \equiv \left\{\boldsymbol{i} = (i_1, i_2, \ldots, i_k) \in \mathbb{N}^k : L_j \leq i_j \leq U_{n,k,j}, \;\; 1 \leq j \leq k\right\}$$

*and*

$$U_{n,k,j} \equiv n - 2(k-j) - 1; \qquad L_j \equiv \begin{cases} 1, & j = 1; \\ 2 + i_{j-1}, & 2 \leq j \leq k. \end{cases} \tag{S47}$$

*Proof.* Let $p \equiv \lfloor n/2 \rfloor$.
**Case I:** If $n$ is odd, then $p = \lfloor n/2\rfloor = \lfloor \frac{n-1}{2}\rfloor$. For Eq. S46 to hold, the right-hand side of Eq. S44 becomes

$$D_n - \gamma_n D_{n-1} = 1 + \sum_{k=1}^{p}(-1)^k \sum_{i\in\mathcal{S}_k^{(n-1)}}\prod_{j=1}^{k}\gamma_{i_j} - \gamma_n\left[1 + \sum_{k=1}^{p}(-1)^k \sum_{i\in\mathcal{S}_k^{(n)}}\prod_{j=1}^{k}\gamma_{i_j}\right]$$

$$= 1 + \sum_{k=1}^{p}(-1)^k\left[\sum_{i\in\mathcal{S}_k^{(n)}}\prod_{j=1}^{k}\gamma_{i_j} + \sum_{i\in\mathcal{S}_{k-1}^{(n-1)}}\gamma_n\prod_{j=1}^{k}\gamma_{i_j}\right] + (-1)^{p+1}\sum_{i\in\mathcal{S}_p^{(n-1)}}\gamma_n\prod_{j=1}^{p}\gamma_{i_j}. \tag{S48}$$

For the claim to be true, it must be that Eq. S48 equates to

$$D_{n+1} = 1 + \sum_{k=1}^{p+1}(-1)^k \sum_{i\in\mathcal{S}_k^{(n+1)}}\prod_{j=1}^{k}\gamma_{i_j}. \tag{S49}$$

We will show this term-by-term. Namely,

$$\begin{aligned} &i: \sum_{i\in\mathcal{S}_k^{(n+1)}} \prod_{j=1}^{k} \gamma_{i_j} = \sum_{i\in\mathcal{S}_k^{(n)}} \prod_{j=1}^{k} \gamma_{i_j} + \sum_{i\in\mathcal{S}_{k-1}^{(n-1)}} \gamma_n \prod_{j=1}^{k} \gamma_{i_j}, \;\; 1\le k\le p, \;\; \text{and;} \\ &ii: \sum_{i\in\mathcal{S}_{p+1}^{(n+1)}} \prod_{j=1}^{p+1} \gamma_{i_j} = \sum_{i\in\mathcal{S}_p^{(n-1)}} \gamma_n \prod_{j=1}^{p} \gamma_{i_j}. \end{aligned} \tag{S50}$$

To account for the addition of the $\gamma_n$ factor in the second term, we define:

$$\tilde{\mathcal{S}}_k^{(n-1)} \equiv \mathcal{S}_{k-1}^{(n-1)} \times \{n\} = \left\{ \boldsymbol{i} \in \mathbb{N}^k : \begin{array}{cc} L_j \le i_j \le U_{n-1,k-1,j}, & 1 \le j \le k-1; \\ i_j = n, & j = k. \end{array} \right\}. \tag{S51}$$

Then we may re-write its corresponding term as:

$$\sum_{i\in\mathcal{S}_{k-1}^{(n-1)}} \gamma_n \prod_{j=1}^{k} \gamma_{i_j} = \sum_{i\in\tilde{\mathcal{S}}_k^{(n-1)}} \prod_{j=1}^{k} \gamma_{i_j}$$

Thus, from Eq. S50, it suffices to show that

$$\begin{aligned} &i: \mathcal{S}_k^{(n+1)} = \mathcal{S}_k^{(n)} \,\dot\cup\, \tilde{\mathcal{S}}_k^{(n-1)}, \;\; 1 \le k \le p, \;\; \text{and;} \\ &ii: \mathcal{S}_{p+1}^{(n+1)} = \tilde{\mathcal{S}}_{p+1}^{(n+1)}. \end{aligned} \tag{S52}$$

In the above, $A \,\dot\cup\, B$ denotes the disjoint union (i.e. $A \cup B$ where $A \cap B = \emptyset$). Corresponding to $\mathcal{S}_k^{(n)}$, put

$$\begin{aligned} &I_j \equiv \left\{ \boldsymbol{i} \cdot e_j : \boldsymbol{i} \in \mathcal{S}_k^{(n)} \right\} = \left\{ i_j : L_j \le i_j \le U_{n,k,j} \right\}; \\ &\tilde{I}_j \equiv \left\{ \boldsymbol{i} \cdot e_j : \boldsymbol{i} \in \tilde{\mathcal{S}}_k^{(n-1)} \right\}. \end{aligned}$$

so that

$$\mathcal{S}_k^{(n)} = \prod_{j=1}^{k} I_j, \qquad \text{and } \tilde{\mathcal{S}}_k^{(n-1)} = \prod_{j=1}^{k} \tilde{I}_j. \tag{S53}$$

Note that since $\tilde{\mathcal{S}}_k^{(n-1)} = \mathcal{S}_{k-1}^{(n-1)} \times \{n\}$, $n = U_{n,k,j} + 1$, and since

$$U_{n-1,k-1,j} = n - 1 - 2(k - 1 - j) - 1 = U_{n,k,j} + 1,$$

this implies that $\tilde{I}_j = \left\{ i_j : L_j \le i_j \le U_{n,k,j} + 1 \right\}$.
To simplify the notation, put $w_j \equiv U_{n,k,j} + 1$. Then

$$\tilde{I}_j = \left\{ \begin{array}{cc} \{w_j\}, & j = k; \\ I_j \,\dot\cup\, \{w_j\}, & 1 \le j \le k-1. \end{array} \right.$$

Clearly $\mathcal{S}_k^{(n)}$, $\tilde{\mathcal{S}}_k^{(n-1)}$ are disjoint since $I_k \cap \tilde{I}_k = \{i_j : L_j \le i_j \le w_j - 1\} \cap \{w_j\} = \emptyset$.
It remains to be shown that their union coincides with $\mathcal{S}_k^{(n+1)}$. We will show this by constructing partial Cartesian products. Toward this end, let $H_j \equiv \left\{ I_j \,\dot\cup\, \{w_j\}, \;\text{ for } 1 \le j \le k \right\}$. Then $H_j = \tilde{I}_j$ for $1 \le j \le k-1$, and since $U_{n+1,k,j} = U_{n,k,j} + 1 = w_j$, we have that

$$\prod_{j=1}^{k} H_j = \mathcal{S}_k^{(n+1)}.$$

Moreover, since $U_{n,k,j} - U_{n,k,j-1} = L_j - L_{j-1} = 2$, it follows that if $i_{j-1} = w_{j-1} = U_{n,k,j-1} + 1$, then

$$i_j \geq L_j = 2 + U_{n,k,j-1} + 1 = U_{n,k,j} + 1 = w_j.$$

Thus, for $\boldsymbol{i} \in \tilde{\mathcal{S}}_k^{(n-1)}$,

$$i_j = w_j \;\Rightarrow\; i_\ell = w_\ell, \;\text{ for }\; j \leq \ell \leq k. \tag{S54}$$

Equivalently,

$$\chi_1 \times \chi_2 \times \cdots \times \chi_k = \emptyset, \;\text{ for } \chi_j \in \Big\{ I_j, \{w_j\} \Big\}$$

whenever

$$\min_{1\leq j\leq k} \Big\{ j : \chi_j = \{w_j\} \Big\} < \max_{1\leq j\leq k} \Big\{ j : \chi_j = I_j \Big\}.$$

In other words, the product is nonempty only when all sets following the earliest appearance of the singleton $\{w_j\}$ are followed by $\{w_{j+1}\}, \{w_{j+2}\}, \ldots$. Now, working backward from the $k^{\text{th}}$product:

$$I_k \,\dot\cup\, \tilde{I}_k = I_k \,\dot\cup\, \{w_k\} = H_k.$$

For the $k-1^{\text{th}}$ product:

$$\begin{aligned} I_{k-1} \times I_k \,\dot\cup\, \tilde{I}_{k-1} \times \tilde{I}_k &= I_{k-1} \times I_k \,\dot\cup\, \Big( I_{k-1} \,\dot\cup\, \{w_{k-1}\} \Big) \times \{w_k\} \\ &= \big( I_{k-1} \times I_k \big) \,\dot\cup\, \big( I_{k-1} \times \{w_k\} \big) \,\dot\cup\, (w_{k-1}, w_k) \\ &= I_{k-1} \times H_k \,\dot\cup\, (w_{k-1}, w_k) \\ &= H_{k-1} \times H_k. \end{aligned}$$

The last equality holds since

$$\begin{aligned} I_j \times H_{j+1} \times \cdots \times H_k = &\Big\{ i_j : L_j \leq i_j \leq w_j - 1 \Big\} \times \Big\{ i_{j+1} : L_{j+1} \leq i_{j+1} \leq w_{j+1} \Big\} \times \cdots \\ &\times \Big\{ i_k : L_k \leq i_k \leq w_k \Big\} \\ = &\, H_j \times \cdots \times H_k \setminus (w_j, \ldots, w_k). \end{aligned}$$

More generally, we now consider

$$\prod_{j=1}^{k} I_j \,\dot\cup\, \prod_{j=1}^{k} \tilde{I}_j$$

We remark that

$$\prod_{j=1}^{k} \tilde{I}_j = \bigcup_{\boldsymbol{\alpha}\in\Lambda} \prod_{j=1}^{k} X_{j,\boldsymbol{\alpha}}, \tag{S55}$$

for

$$\Lambda_k \equiv \Big\{ \boldsymbol{\alpha} = (\alpha_1, \ldots, \alpha_k) \in \{0,1\}^k : \alpha_k = 1 \Big\},$$

and

$$X_{j,\boldsymbol{\alpha}} \equiv \begin{cases} \{w_j\}, & \alpha_j = 1; \\ I_j, & \alpha_j = 0. \end{cases}$$

By the argument in Eq. S54, we have that

$$\bigcup_{\boldsymbol{\alpha}\in\Lambda} \prod_{j=1}^{k} X_{j,\boldsymbol{\alpha}} = \bigcup_{z=1}^{k} \prod_{j=1}^{k} X_{j,\boldsymbol{\beta}_z}, \tag{S56}$$

for

$$\boldsymbol{\beta}_z = \left\{ (\boldsymbol{\beta}_{z,1}, \ldots, \boldsymbol{\beta}_{z,k}) \in \{0,1\}^k : \begin{array}{ll} \boldsymbol{\beta}_{z,j} = 1, & j > k - z; \\ \boldsymbol{\beta}_{z,j} = 0, & \text{otherwise.} \end{array} \right\}.$$

Thus, by Eqs. S53,S56 this gives for the desired quantity,

$$\begin{aligned}
\mathcal{S}_k^{(n)} \dot\cup \tilde{S}_k^{(n-1)} &= \prod_{j=1}^{k} I_j \dot\cup \bigcup_{z=1}^{k} \prod_{j=1}^{k} X_{j,\boldsymbol{\beta}_z} \\
&= \bigcup_{z=0}^{k} \prod_{j=1}^{k} X_{j,\boldsymbol{\beta}_z} \\
&= \bigcup_{z=2}^{k} \prod_{j=1}^{k} X_{j,\boldsymbol{\beta}_z} \dot\cup \prod_{j=1}^{k} X_{j,\boldsymbol{\beta}_0} \dot\cup \prod_{j=1}^{k} X_{j,\boldsymbol{\beta}_1} \\
&= \bigcup_{z=3}^{k} \prod_{j=1}^{k} X_{j,\boldsymbol{\beta}_z} \dot\cup \prod_{j=1}^{k-1} I_j \times H_k \dot\cup \prod_{j=1}^{k} X_{j,\boldsymbol{\beta}_2} \\
&\vdots \\
&= \prod_{j=1}^{k} X_{j,\boldsymbol{\beta}_k} \dot\cup I_1 \times H_2 \times \cdots \times H_k \\
&= \prod_{j=1}^{k} H_j = \mathcal{S}_k^{(n+1)},
\end{aligned}$$

proving Condition *ii* of Eq. S52.

For Condition *i*, by definition,

$$\tilde{\mathcal{S}}_{p+1}^{(n-1)} = \left\{ \boldsymbol{i} \in \mathbb{N}^{p+1} : i_{p+1} = n;\ L_j \le i_j \le U_{n-1,p+1,j},\ \ 1 \le j < p+1 \right\}.$$

However,

$$U_{n+1,p+1,j} = n + 1 - 2(p + 1 - j) - 1 = n - 1 - 2(p - j) - 1 = U_{n-1,p,j},$$

and

$$i_{p+1} = n = n + 1 - 2[p + 1 - (p + 1)] - 1 = U_{n+1,p+1,p+1}.$$

Thus, in this particular case,

$$\tilde{\mathcal{S}}_{p+1}^{(n-1)} = \left\{ \boldsymbol{i} \in \mathbb{N}^{p+1} : L_j \le i_j \le U_{n+1,p+1,j},\ \ 1 \le j \le p+1 \right\} = \mathcal{S}_{p+1}^{(n)}.$$

Proving Condition *ii*.

**Case II:** If $n$ is even, then $p = \lfloor n/2 \rfloor = \lfloor \frac{n+1}{2} \rfloor$. In this case, Eqs. S48-S49 become

$$D_{n+1} = 1 + \sum_{k=1}^{p} (-1)^k \sum_{i \in \mathcal{S}_k^{(n+1)}} \prod_{j=1}^{k} \gamma_{i_j}$$

and

$$D_n - \gamma_n D_{n-1} = 1 + \sum_{k=1}^{p} (-1)^k \sum_{i \in \mathcal{S}_k^{(n)}} \prod_{j=1}^{k} \gamma_{i_j} - \gamma_n \left[ 1 + \sum_{k=1}^{p-1} (-1)^k \sum_{i \in \mathcal{S}_k^{(n-1)}} \prod_{j=1}^{k} \gamma_{i_j} \right].$$

Term-by-term, this becomes

$$\sum_{i\in\mathcal{S}_k^{(n+1)}}\prod_{j=1}^{k}\gamma_{i_j}=\sum_{i\in\mathcal{S}_k^{(n)}}\prod_{j=1}^{k}\gamma_{i_j}+\sum_{i\in\mathcal{S}_{k-1}^{(n-1)}}\gamma_n\prod_{j=1}^{k}\gamma_{i_j},\ \ 1\le k\le p,$$

Which follows from directly as in Condition $i$ of Eq. S50

□

The general case, $d\ge 0$, follows from Eq. S46, with $D_d = D_{d+1} = 1$, and, for $n\ge 2$:

$$D_{d+n}=1+\sum_{k=1}^{\lfloor\frac{d+n}{2}\rfloor}(-1)^k\sum_{i\in{}^d\mathcal{S}_k^{(n)}}\prod_{j=1}^{k}\gamma_{i_j},\tag{S57}$$

where

$${}^d\mathcal{S}_k^{(n)}\equiv\Big\{\boldsymbol{i}=(i_1,i_2,\ldots,i_k)\in\mathbb{N}^k : L_j+d\le i_j\le U_{n,k,j}+d,\ \ 1\le j\le k\Big\}.$$

From this, it follows that

$$\mathbb{P}\left({}^dE_{n+1,n}\right)=\frac{D_n}{D_{n-1}}\nu_n,\quad n\ge d+1.$$

Thus, solutions may be solved analytically or calculated recursively for:

$$\mathbb{P}\left({}^dE_{n,d+1}\right)=\prod_{j=d+2}^{n-1}\frac{D_j}{D_{j-1}}\nu_j=\frac{D_{n-1}}{D_d}\prod_{j=d+1}^{n-1}\nu_j,\quad n\ge d+1,$$

and

$$\mathbb{P}\left({}^dE_{n,m}\right)=\frac{D_{n-1}}{D_{m-1}}\prod_{j=m}^{n-1}\nu_j,\quad n>m>d.\tag{S58}$$

##### S8.1.1 Linear Fractional Transform

In the previous section S8.1, we derived an analytical representation for the conditional hitting probabilities for inhibitory signal size and recursively calculated them. However, evaluating the closed from of the probabilities is computationally intensive as we need to evaluate the sequence of real numbers, $D_n$, given in (S45). To that extent, we apply an interesting fact from complex analysis to provide a convenient way to compute the hitting probabilities. Consider the following linear fractional transformation from the complex numbers $\mathbb{C}$ to itself defined by

$$f(z)=\frac{az+b}{cz+e},\qquad \text{with } a,b,c,e,z\in\mathbb{C}\text{ s.t. } ac-be\ne 0$$

We state the following facts regarding this transformation:

- Linear fractional transformations form a group under the action of compositions (composition of two linear fractional transform is another linear fractional transform),
- The function mapping the space of $2\times 2$ invertible matrices ($GL_2(\mathbb{C})$) to the space of linear fractional transformations is surjective: any linear fractional transformation comes from a $2\times 2$ invertible matrix. For example, $f$ defined earlier can be identified by $\begin{bmatrix}a & b\\ c & e\end{bmatrix}$ matrix action on $z$,

- $f$ is an map from the real line to itself $\iff a, b, c, e \in \mathbb{R}$.

The first two facts imply that given two linear fractional transformations $f, g$ with their associated $2 \times 2$ invertible matrices $A, B$, then $f \circ g$ is also a linear fractional transformation with its associated matrix given by $A \times B$. From the previous section, we have

$$\begin{aligned}\mathbb{P}\left({}^{d}E_{n+1,n}\right) &= \left(\frac{\lambda_n}{\lambda_n+\mu_n}\right) + \mathbb{P}\left({}^{d}E_{n+1,n-1}\right)\left(\frac{\mu_n}{\lambda_n+\mu_n}\right)\\ &= \nu_n + \eta_n\,\mathbb{P}\left({}^{d}E_{n,n-1}\right)\,\mathbb{P}\left({}^{d}E_{n+1,n}\right),\end{aligned}$$

where we have defined

$$\nu_n \equiv \frac{\lambda_n}{\lambda_n+\mu_n}; \qquad \eta_n \equiv \frac{\mu_n}{\mu_n+\lambda_n}.$$

Note that we have defined $\eta_n$ differently from Eq. (S42) in the previous section. In addition, we have hidden the immunomodulatory dependency from the rate functions. Assume that calculation is for a constant initial immunomodulatory signal, $M$. This yields the recursive relation

$$\mathbb{P}\left({}^{d}E_{n+1,n}\right) = \frac{\nu_n}{1-\eta_n\,\mathbb{P}\left({}^{d}E_{n,n-1}\right)}, \quad \text{with } \mathbb{P}\left({}^{d}E_{d+2,d+1}\right) = \nu_{d+1}.$$

Let $x_n = \mathbb{P}\left({}^{d}E_{n+1,n}\right)$ and $f_n$ the linear fractional transformation associated with the matrix $J_n = \begin{bmatrix} 0 & \nu_n \\ -\eta_n & 1 \end{bmatrix}$, then the above recursion can be written as applications of linear fractional transformations from the real line to the real line (as all entries in $J_n$ are real):

$$x_n = f_n(x_{n-1}) = \frac{\nu_n}{1-\eta_n x_{n-1}}, \quad \text{with } x_{d+1} = \nu_{d+1}.$$

We exploit this recursive relationship and the boundary condition to write our equations as a composition of linear fractional transformations, namely,

$$\begin{aligned}x_n &= f_n(x_{n-1})\\ &= f_n \circ f_{n-1}(x_{n-2})\\ &= f_n \circ \cdots \circ f_{d+2}(\nu_{d+1}).\end{aligned}$$

Now, each LFT $f_n$ has its associated matrix $J_n$. As such, via the group property of LFT, we have that $f_n \circ \cdots \circ f_{d+2}$ is also another LFT with its associated matrix given by

$$\prod_{i=n}^{d+2} J_i = J_n \times J_{n-1} \times \cdots \times J_{d+2}.$$

This provides an computationally tractable method to evaluate the hitting probability by viewing it as a product of matrix products as opposed to the analytical form in Eq. (S58). Namely,

$$\mathbb{P}\left({}^{d}E_{n+1,n}\right) = \frac{a\nu_{d+1}+b}{c\nu_{d+1}+e} \quad \text{with } \begin{bmatrix} a & b \\ c & e \end{bmatrix} \triangleq \prod_{i=n}^{d+2} \begin{bmatrix} 0 & \nu_i \\ -\eta_i & 1 \end{bmatrix}. \tag{S59}$$

We use this method of computation to look into step up probabilities and escape probabilities for the passive immunomodulation case under a constant immunomodulatory landscape and a high per cell killing rate.

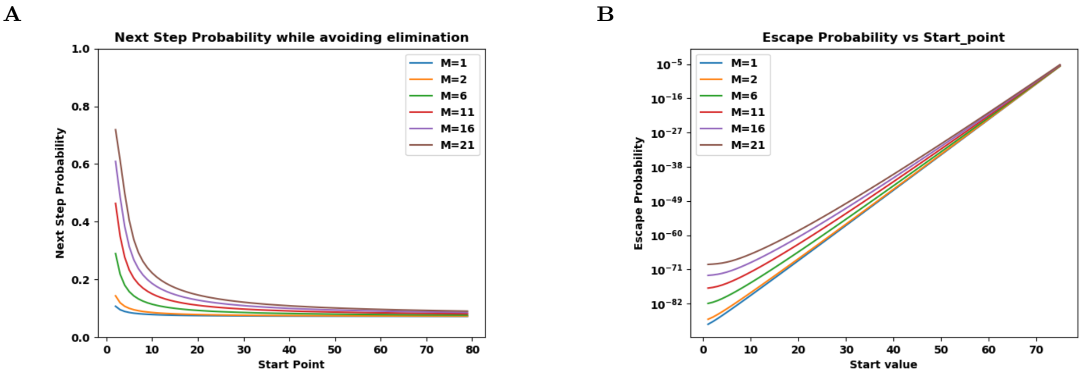


Figure S6: Next step probabilities and escape probabilities for passive immunomodulation via LFT under a constant immunomodulatory landscape **A:** Probability of a unit increase in tumor size without being eliminated first for passive immunomodulatory function **B:** Escape probability (in log-scale) vs initial start value using the results for next step probabilities for passive immunomodulatory function. ($r = 0.05, \delta = 0.7, N = 120, \alpha = 0.1$, and $\beta = 0.18$.)

#### S8.2 Application of dynamic immunomodulation

In this section, we apply the above framework to address cases where the number of immunomodulatory cells, $M$, may depend on the dynamics of the cancer population. We assume that changes in $M$ may be characterized by two features: first, the cancer population sizes that trigger a change in the immunomodulatory population, and second, the extent of that change. Toward this end, we characterize the dynamics using two sequences: the first sequence $\{\delta_1, \delta_2, \dots\}$, $\delta_i \geq 0$ is a collection of cancer population sizes at which the value of $M$ changes. $\{M_1, M_2, \dots\}$, $M_i \geq 0$ represents these values, so that $M = M_i$ whenever $\delta_i \leq X < \delta_{i+1}$. In this framework, the static case may be viewed as the limit where $\min_i |\delta_i - \delta_{i-1}| \to \infty$.

Here, the birth and death rates used to calculate the relevant conditional probabilities are functions of both $n$ and $M$, given by $\lambda_{n,M}$ and $\mu_{n,M}$, respectively. We assume that the cancer population starts at initial size $n_i$ with $M_i$ suppressors. While this framework can handle any generalized suppressor cell dynamics given by $\{M_i\}$ and $\{\delta_i\}$, we consider several particular cases below:

In the analysis below, we assume that $M_i$ is an increasing non-negative sequence of immunomodulatory cells, so that larger (resp. smaller) cancer population sizes result in additional (resp. diminished) immunomodulation. This assumption is consistent with the fact that more cancer cells impair cancer killing through the presence of dynamical immune inhibitory populations. However, this same framework can be applied to arbitrary sequences as well. We subsequently consider two cases: The first, which we refer to as *accumulating immunomodulation*, describes the dynamics assuming that immunomodulatory cells only change once the population achieves a new maximal size. This assumption is relaxed under *reversible immunomodulation.*

##### S8.2.1 Accumulating immunomodulation

Here, we assume that immunoinhibitory signals depend on the cancer population size and, once present, exist permanently. This implies that $M(t)$ is increasing in time, and represents an accumulation of inhibitory signal that increases once a certain cancer population size threshold is achieved. This is particularly relevant in, for example, a model of stroma and recruitment of Tregs inducing CD8+ T cell tolerance, or activation of cancer associated fibroblasts.

The accumulation of inhibitory signal is monotonically increasing in time so that once a certain cancer population size threshold is achieved, additional suppressive cells are recruited permanently. Using the framework above, this means that the number of suppressor cells is related to the maximal threshold size currently achieved in the $\delta$ sequence:

$$M(t) = M_{I(t)}, \qquad I(t) = \max\left\{ i \in \mathbb{Z} : \delta_i \leq \max_{0\leq\tau\leq t} X(\tau) \right\}. \tag{S60}$$

We assume that $\{\delta_i\}$ is an arithmetic sequence with common difference $\Delta \equiv \delta_{i+1} - \delta_i$ for $i \geq k^-$. The only exceptional interval is $\Delta_0 = \delta_{k^-+1} - 1$. For ease of calculation, we will further assume that $\Delta \geq 1$ is an odd integer, and that the process begins at a point $x_0 \equiv X(t=0) > 0$ halfway between two partition elements $\delta_0$ and $\delta_1$. Thus $\delta_0 = x_0 + (\Delta - 1)/2$. We must also identify the limit indices for which the partition covers all states left of $x_0$. Toward that end, we let $k^-$ be the largest partition element so that $\delta_{k^-} \leq 1$. In this case, $\delta_{k^-} \leq 1 < \delta_{k^-+1}$. This lower index can be similarly represented by the extent to which $\delta_{k^-}$ is below $\delta_0$. Namely, that $k^- = -\lceil(\delta_0 - 1)/\Delta\rceil$. Using these indices, we may calculate via iterating the recursive relation given by Eq. S43, and using the notation ${}^dE^M_{n,m}$ to denote the probability of reaching $n$ starting from $m$ conditional on avoiding $d$ with immunomodulation $M$, the sequences describing the probabilities relevant for each level of $M_I$ are given by

$$\begin{aligned}
S_{k^-} &= \left\{ \mathbb{P}\left({}^0E^{M_{k^-}}_{2,1}\right), \mathbb{P}\left({}^0E^{M_{k^-}}_{3,2}\right), \ldots, \mathbb{P}\left({}^0_{M_{k^-}}E_{\delta_{k^-+1},\delta_{k^-+1}-1}\right) \right\}, \\
S_{k^-+1} &= \left\{ \mathbb{P}\left({}^0E^{M_{k^-+1}}_{\delta_{k^-+1,k^-}}\right), \mathbb{P}\left({}^0E^{M_{k^-+1}}_{\delta_{k^-+2,k^-+1}}\right), \ldots, \mathbb{P}\left({}^0E^{M_{k^-+1}}_{\delta_{k^-+2},\delta_{k^-+2}-1}\right) \right\}, \\
&\vdots \\
S_k &= \left\{ \mathbb{P}\left({}^0E^{M_k}_{\delta_{k+1,k}}\right), \mathbb{P}\left({}^0E^{M_k}_{\delta_{k+2,k+1}}\right), \ldots, \mathbb{P}\left({}^0E^{M_k}_{\delta_{k+1,},\delta_{k+1}-1}\right) \right\}.
\end{aligned} \tag{S61}$$

From this, we may calculate the corresponding probabilities

$$\begin{aligned}
\varphi_{\delta_{k^-},1} &\equiv \mathbb{P}\left({}^0E^{M^-_k}_{\delta_{k^-+1},1}\right) = \prod_{j=1}^{\Delta_0-1} \mathbb{P}\left({}^0E^{M_{k^-}}_{j+1,j}\right), \\
\varphi_{\delta_{k^-+1},\delta_{k^-}} &\equiv \mathbb{P}\left({}^0E^{M^-_k+1}_{\delta_{k^-+2},\delta_{k^-+1}}\right) = \prod_{j=1}^{\Delta-1} \mathbb{P}\left({}^0E^{M_{k^-+1}}_{\delta_{k^-}+j+1,\delta_{k^-}+j}\right), \\
&\vdots \\
\varphi_{\delta_{k+1},\delta_k} &\equiv \mathbb{P}\left({}^0E^{M_k}_{\delta_k+1,\delta_k}\right) = \prod_{j=1}^{\Delta-1} \mathbb{P}\left({}^0E^{M_k}_{\delta_k+j+1,\delta_k+j}\right).
\end{aligned} \tag{S62}$$

From this, the escape probability can be calculated as

$$P_{esc} = \varphi_{\delta_{k^-},1} \prod_{k=k^-+1}^{\infty} \varphi_{\delta_k,\delta_{k-1}}. \tag{S63}$$

Alternately, via the linear fractional transform approach, for any initial size $m$, the escape probability can be calculated as

$$P_{esc} = \prod_{n=m}^{\infty} \mathbb{P}\left({}^0E_{n+1,n} \,:\, M = M_n\right) \tag{S64}$$

where $M_n$ is the immodulation signal size with respect to tumor size $n$. From our accumulatory landscape assumption, at size $n$, the immodulatory signal does not change unless the tumor size increases. As such, the tumor fluctuates with a constant immunomodulatory signal, $M = M_n$, and increasing only when the tumor grows above $n$. Thus, we can use the results from Eq. (S59) with $d = 0$ to calculate $\mathbb{P}\left({}^0E_{n+1,n} \,:\, M = M_n\right)$. The infinite sum in Eq. (S64), by virtue of the increasing nature of $M_n$, accounts for accumulation.

Plots of accumulating immunomodulation transition probabilities and overall escape probabilities using the results of Sec S8.1 is given by Fig. S7 whereas the plots of the corresponding stochastic trajectories and immunomodulation values given in Fig. S9. Finally, the plots of the accumulation immunomodulation transition and escape probabilities using linear fractional transform as shown in Sec S8.1.1 is given in Figs S10 and S11.

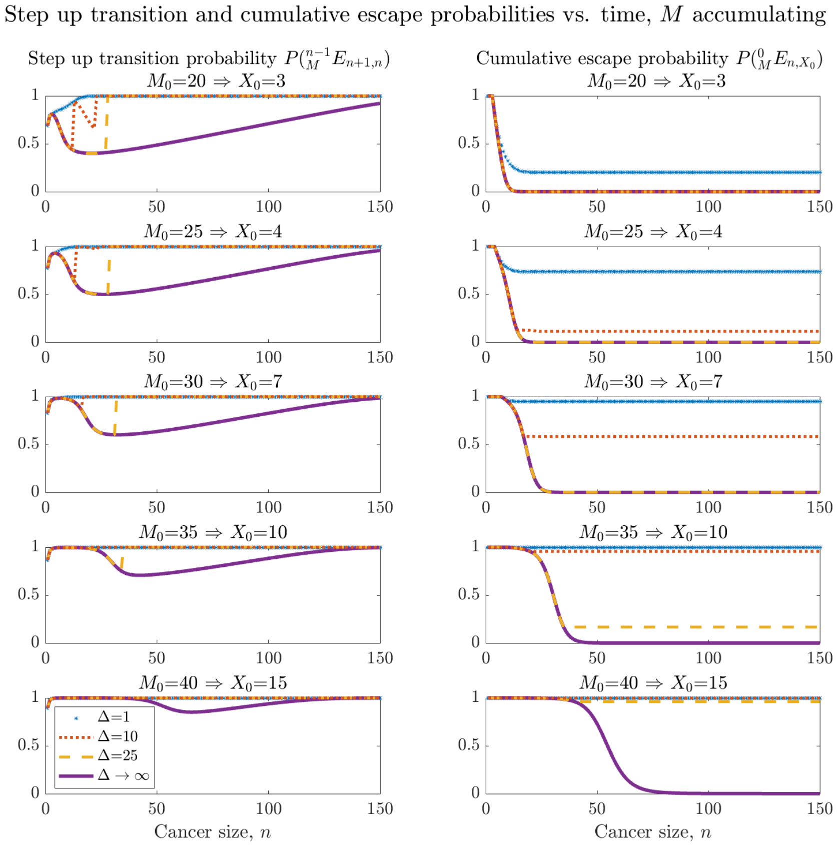


Figure S7: Accumulating immunomodulation step up transition and cumulative escape probabilities. Step up transition probabilities (left column) and cumulative escape probabilities (right column) are calculated iteratively according to Sec. S8.2.1 and plotted for various difference $\Delta$ values. Plots are given for various immune inhibition (minimal immune inhibition on top; maximal immune inhibition on bottom. In all cases, $f$ is used according to linear inhibition case in Eq. (S37) with $r = 0.005$, $d = 1$, $\alpha = \beta = 1$).

##### S8.2.2 Reversible immunomodulation

In contrast to accumulating immunomodulation, reversible immunomodulation allows for increases and decreases in $M$ based on cancer population growth and reduction, respectively. This application may be particularly relevant for modeling, for example, the reversible impact of myeloid-derived stem cells. The number of suppressor cells is related to the current threshold size:

$$M(t) = M_{I(t)}, \qquad I(t) = \max\{i \in \mathbb{Z} : \delta_i \leq X(t)\}. \tag{S65}$$

We may proceed similarly in this case, except that $M = M_0$ for all state values below and including $\delta_0$.

Namely,

$$\varphi_{\delta_{k^-},1} = \prod_{j=1}^{\Delta_0 - 1} \mathbb{P}\left({}^0E^{M_0}_{j+1,j}\right),$$

$$\vdots$$

$$\varphi_{\delta_0,\delta_{-1}} = \prod_{j=1}^{\Delta - 1} \mathbb{P}\left({}^0E^{M_0}_{\delta_{-1}+j+1,\delta_{-1}+j}\right),$$

$$\varphi_{\delta_1,\delta_0} = \prod_{j=1}^{\Delta - 1} \mathbb{P}\left({}^0E^{M_1}_{\delta_0+j+1,\delta_0+j}\right),$$

$$\vdots$$

$$\varphi_{\delta_{k+1},\delta_k} = \prod_{j=1}^{\Delta - 1} \mathbb{P}\left({}^0E^{M_k}_{\delta_k+j+1,\delta_k+j}\right). \tag{S66}$$

The escape probability is then calculated identically, so that either $\delta_0 = x_0$ or $\delta_0 < x_0 < \delta_1$. In order to iteratively describe the escape probability, we must first identify the $\delta$ indices around $x = 0$.

$$\mathbb{P}\left({}^0_{M_{k^-+1}}E_{\delta_{k^-+1}+1,\delta_{k^-}}\right) \tag{S67}$$

Alternatively, by linear fractional transform, we can again view this as matrix multiplication. From Eq. (S64), the infinite sum accounts for the accumulatory aspect of the landscape. The accumulatory landscape assumption further allowed us to view the immunomodulatory signal as constant when calculating the elements of the infinite sum. This is not true for the reversible landscape. Namely, let $n, k$ be such that $\delta_k \leq n < \delta_{k+1}$ then

$$\mathbb{P}\left({}^0E_{n+1,n} \,:\, M = M_n\right) = \frac{a\nu_1 + b}{c\nu_1 + e} \quad \text{with} \tag{S68}$$

$$\begin{bmatrix} a & b \\ c & e \end{bmatrix} \triangleq \prod_{i=n}^{\delta_k} \begin{bmatrix} 0 & \nu_i \\ -\eta_i & 1 \end{bmatrix}_{M_{\delta_k}} \times \prod_{j=k-1}^{1} \left( \prod_{i=\delta_{j+1}}^{\delta_j} \begin{bmatrix} 0 & \nu_i \\ -\eta_i & 1 \end{bmatrix}_{M_{\delta_j}} \right) \times \prod_{i=\delta_1}^{2} \begin{bmatrix} 0 & \nu_i \\ -\eta_i & 1 \end{bmatrix}_{M_{\delta_0}}, \tag{S69}$$

where $[\cdot]_x$ means that the matrix rates are evaluated with immunomodulatory parameter $x$. Plots of reversible immunomodulation transition probabilities and overall escape probabilities using the iterative function from Sec S8.1 is given by Fig. S8. In addition, the corresponding stochastic trajectories and immunomodulation values are illustrated in Fig. S9. Moreover, the plots of the reversible immunomodulation transition and escape probabilities using the results of Sec S8.1.1 is given in Figs S10 and S11.

##### S8.2.3 Applications

In both cases, we will illustrate the behavior of dynamical immunomodulation by focus our calculations on the case where both $\{\delta_i\}$ and $\{M_i\}$ are arithmetic sequences with common difference $\Delta > 0$. The accumulating immunomodulation model appears to fit the example where $M$ represents MHC-I expression of a cancer population. Interestingly, MHC high ($M$ low) is quickly eliminated if doesn't move to MHC-I medium. There, the population may stay, perhaps for awhile if the barrier to total MHC-I depletion is large (represented by large $\Delta_i$ in this regime, before MHC-I depleted escape.

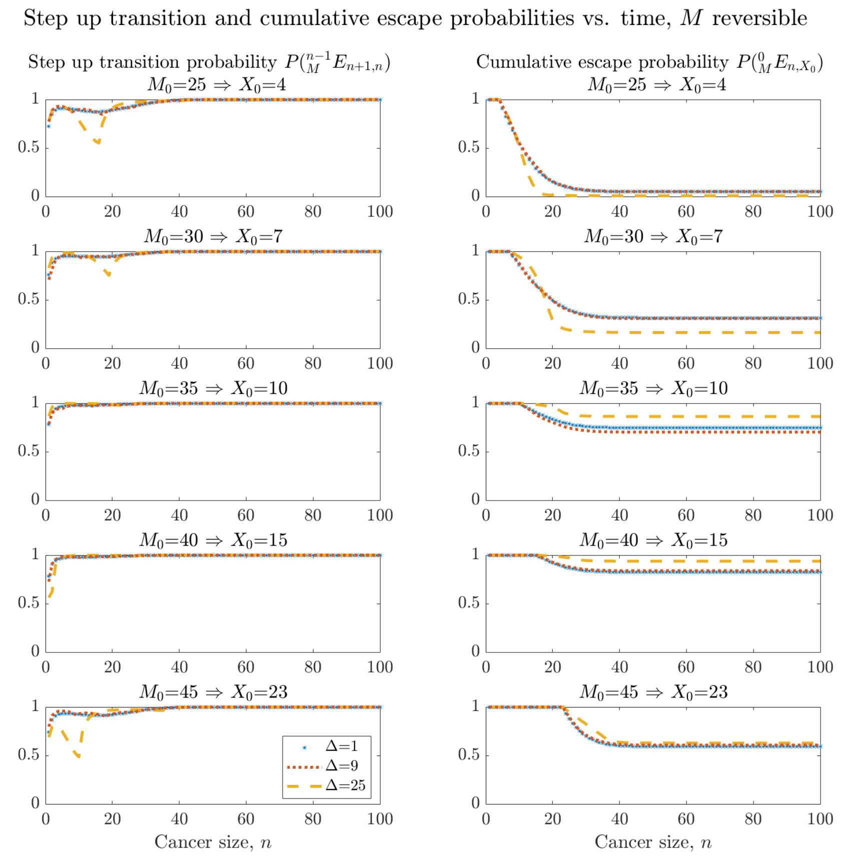


Figure S8: Reversible immunomodulation step up transition and cumulative escape probabilities. Step up transition probabilities (left column) and cumulative escape probabilities (right column) are calculated iteratively according to Sec. S8.2.2 and plotted for various difference $\Delta$ values. Plots are given for various immune inhibition (minimal immune inhibition on top; maximal immune inhibition on bottom. In all cases, $f$ is used according to linear inhibition case in Eq. (S37) with $r = 0.005$, $d = 1$, $\alpha = \beta = 1$).

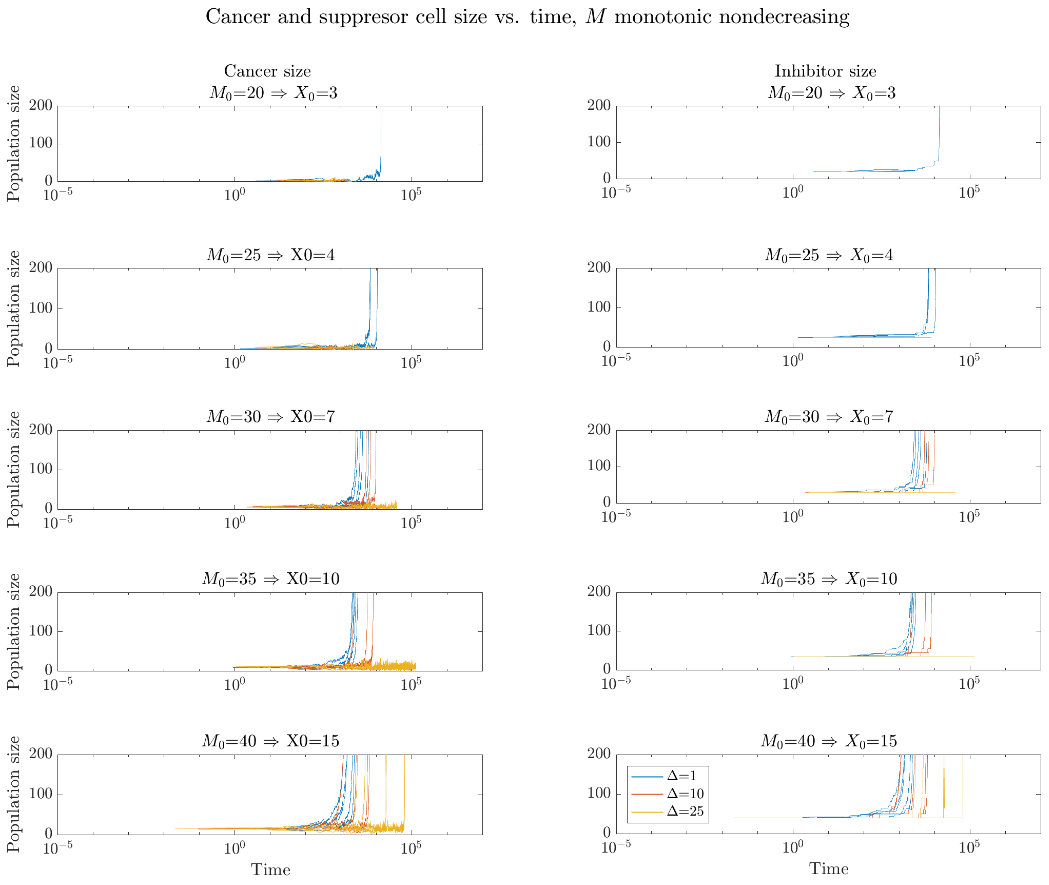


Figure S9: Tumor and immune inhibition dynamics under accumulating immunomodulation. Tumor size $X(t)$ (left column) and accumulating inhibition size $M(t)$ (right column) are plotted for as a function of increasing initial inhibition size $M_0$ (top rows to bottom rows). For each assumed value of difference $\Delta$, a total of five iterations are reported. In all cases, $f$ is used according to linear inhibition case in Eq. (S37) with $r = 0.005$, $d = 1$, $\alpha = \beta = 1$.

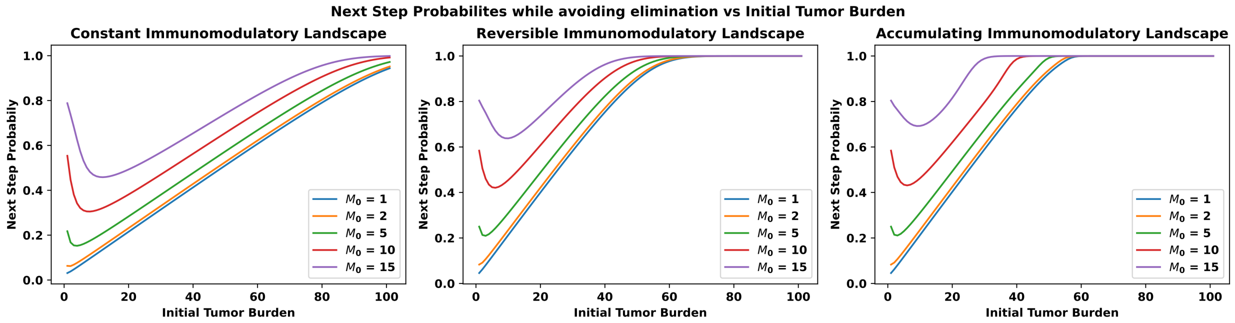


Figure S10: Probability of a unit increase in tumor size without being eliminated first via Linear fractional transform for constant, reversible and accumulatory landscapes respectively. In all cases, $f$ is used according to linear inhibition case in Eq. (S37) with $M_0$ as the initial immunomodulatory signal and $r = 0.05, d = 0.9, N = 150, \alpha = 0.1$, and $\beta = 0.18$.. $\Delta_n = 1$, $\Delta_M = 0.5$ where every $\Delta_n$ change (increase for accumulating) in tumor size results in $\Delta_M$ change (increase resp.) in the immunomodulatory signal.

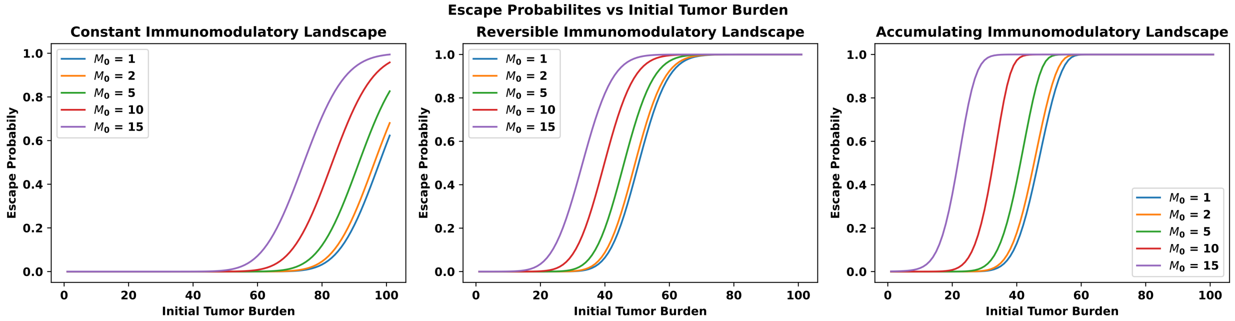


Figure S11: Escape probability vs initial start value using the results for next step probabilities under constant, and accumulatory landscapes respectively. In all cases, $f$ is used according to linear inhibition case in Eq. (S37) with $M_0$ as the initial immunomodulatory signal and $r = 0.05, d = 0.9, N = 150, \alpha = 0.1$, and $\beta = 0.18$.. $\Delta_n = 1$, $\Delta_M = 0.5$ where every $\Delta_n$ change (increase for accumulating) in tumor size results in $\Delta_M$ change (increase resp.) in the immunomodulatory signal.

### S9 Application to Clinical Incidence Data

Lastly, we apply these findings to cancer incidence data for patients with breast and bladder cancer. In the absence of clinical micro-metastasis longitudinal tumor trajectory data, we abstract the notion of escape to each individual progression. As such, the data consists of incidence record from diagnosis to first progression, first to second and so on. By doing so, we obtain the distribution for the "time to escape" for each progression. We fit the model to breast and bladder cancer clinical incidence data with this interpretation and we estimate the immunomodulation parameters for each event interval. The breast cancer data were obtained from an online repository [37] associated with the breast cancer forecasting work by Newton et al. [38] which utilized clinical data from Memorial Sloan Kettering Cancer Center (MSKCC) and MD Anderson Cancer Center (MDACC). Similarly, the bladder cancer data were sourced from an online repository [39] associated with the modeling work by Hasnain et al. [40] which uses clinical data from USC Institute of Urology.

#### S9.1 Data Acquisition

To analyze the dynamics of metastatic progression in early-stage breast and bladder cancer, we utilized longitudinal datasets from the MSKCC, MDACC, and USC Institute of Urology. These datasets provided detailed clinical and demographic information for 3,505 bladder cancer patients and 4,181 breast cancer patients.

For bladder cancer, we focused on a dataset from the USC Institute of Urology, which contained comprehensive data on 3,505 patients. These patients were monitored from their cystectomy through successive stages of metastatic progression: cystectomy to 1st progression, 1st to 2nd progression, 2nd to 3rd progression, and 3rd to 4th progression. The dataset included detailed patient information such as age, sex, primary histology, Charlson Comorbidity Index, neoadjuvant and adjuvant chemotherapy, lymphovascular invasion, pathologic subgroup, and decade of diagnosis.

For breast cancer, we utilized two longitudinal datasets. The first dataset included 446 early-stage breast cancer patients diagnosed between 1975 and 2009 at MSKCC, with follow-up extending until 2016. The second dataset comprised 3,735 early-stage breast cancer patients diagnosed at MDACC, with follow-up extending until 2013. In both datasets, patients were followed from their initial diagnosis throughout their metastatic progression and treatment schedules. The stages of progression recorded were diagnosis to 1st progression, 1st to 2nd progression, 2nd to 3rd progression, and 3rd to 4th progression. At the time of diagnosis, none of the patients presented with metastases; however, all eventually developed metastatic

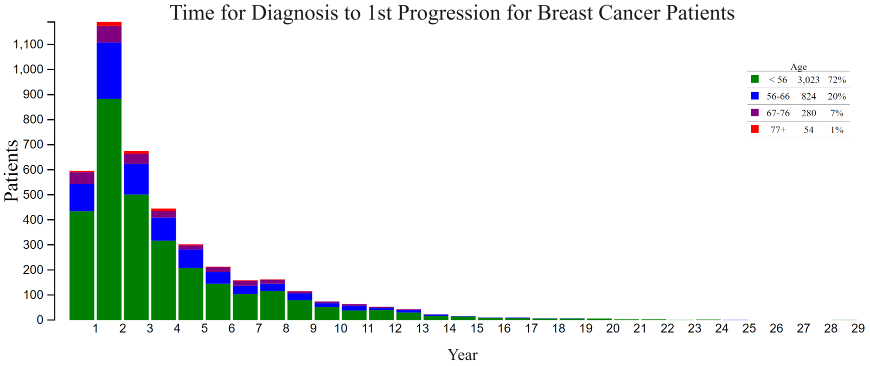


Figure S12: Time from diagnosis to first progression for breast cancer patients, stratified by age groups. The histogram shows the number of patients progressing at each year following diagnosis. The data is categorized into four age groups: under 56 years (green), 56 – 66 years (blue), 67 – 76 years (purple), and over 77 years (red). This clinical data is adapted from [37]

disease. As of August 2013, 273 MDACC patients were deceased, while by July 2016, 2,628 MSKCC patients had succumbed to the disease.

#### S9.2 Data Preprocessing

For each progression stage—diagnosis to 1st progression, 1st to 2nd progression, 2nd to 3rd progression, and 3rd to 4th progression for breast cancer, and cystectomy to 1st progression, 1st to 2nd progression, 2nd to 3rd progression, and 3rd to 4th progression for bladder cancer—we recorded the patient population by year, capturing the age range at each progression point. We then calculated the cumulative distribution function (CDF) for each stage, which represents the cumulative probability of patients progressing to the next stage over time. These CDFs provide a comprehensive overview of the time intervals between successive progression events, offering valuable insights into disease dynamics.

Fig. 6A in the main text illustrates the CDFs for each progression stage in bladder cancer, highlighting the temporal patterns associated with each metastatic event. The CDF for the cystectomy to 1st progression stage shows a more gradual increase, indicating greater variability in progression times. Subsequent progression stages exhibit sharper rises and quicker plateaus, reflecting more rapid transitions. Notably, the CDFs for bladder cancer stages increase more quickly overall compared to breast cancer, suggesting a more aggressive progression of the disease. This analysis provides crucial insights into the metastatic progression timeline in early-stage bladder cancer patients, aiding in the understanding of disease dynamics and potentially guiding treatment strategies.

Similarly, Fig. 6B in the main text illustrates the CDFs for each progression stage in breast cancer. The CDF for the diagnosis to 1st progression stage shows a more gradual increase, reflecting greater variability in progression times among patients. In contrast, the CDFs for subsequent progressions rise sharply and plateau quickly, indicating more rapid transitions between these stages. This analysis offers valuable insights into the metastatic progression timeline in early-stage breast cancer patients, contributing to a better understanding of disease dynamics and potentially informing treatment strategies.

#### S9.3 Fitted Results

In this study, we investigate the dynamics of tumor cell population growth and its modulation by immune suppressor cells. The net growth rate, $\xi(x)$, is a critical determinant of tumor progression, as described by Eq.(10). This growth rate encapsulates both the intrinsic proliferative potential of the tumor cells and the inhibitory effects mediated by the suppressor cell population. By modeling $\xi(x)$, we can estimate the progression time for various tumor cell populations, which is inversely related to $\xi(x)$. This inverse relationship provides a robust framework for understanding the impact of suppressor cell dynamics and other key parameters on tumor growth.

To further analyze the temporal evolution of the tumor cell population, we employ the differential equation $\frac{dx}{dt} = \xi(x)$, which allows us to derive the CDF of the tumor cell population over time. This probabilistic framework offers a comprehensive understanding of tumor progression and yields significant insights for potential therapeutic interventions. By fitting this model to empirical data from large-cohort studies, we can assess how well different immunomodulation scenarios—such as constant, reversible, and accumulating immune suppression—explain the observed progression patterns. This fitting process not only helps in validating the model but also in predicting the extent of immune impairment during different stages of tumor progression.

To validate our model, we calculated the CDF for each progression scenario under constant, reversible, and accumulating immunomodulation. We derived the CDF from clinical data adapted from [37] and [39] for each progression stage and then fitted this data to these immunomodulation landscapes. Specifically, we compared the empirical distributions of inter-progression times to our theoretical predictions by adjusting key parameters such as $\alpha$, $\delta$, and the immunomodulation extent $M$ for each cancer type. We then calculated the mean squared error (MSE) for each progression stage by comparing our model estimates with the clinical data, and subsequently averaged these values. Our analysis revealed that the reversible case consistently exhibited a lower average MSE compared to the constant and accumulating cases, indicating a superior fit

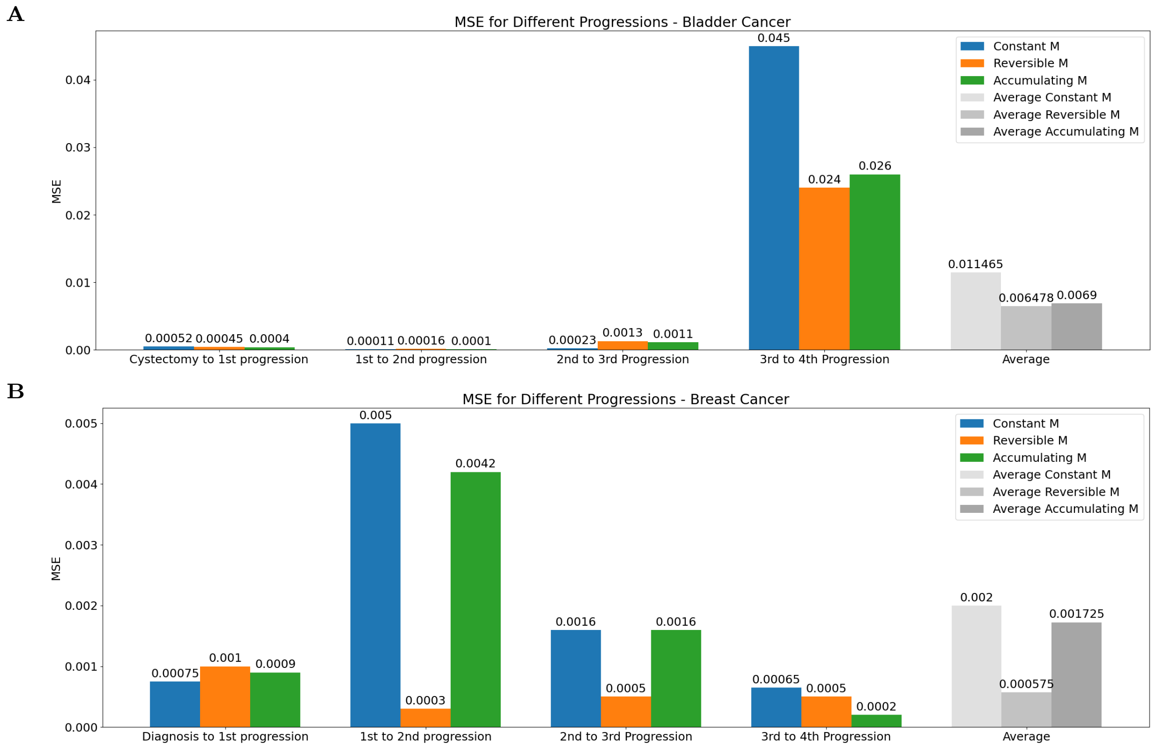


Figure S13: Comparison of mean squared error (MSE) for different progressions stages under constant, reversible, and accumulating immunomodulation for (A) bladder and (B) breast cancer.

and greater accuracy of the reversible immunomodulation. Notably, while all three models were effective in capturing early progression events, the reversible model provided the best fit across the entire progression timeline, particularly in breast cancer. In the following section, we will discuss these findings in detail and explore their implications for understanding cancer progression and optimizing treatment strategies.

##### S9.3.1 Constant Immunomodulatory Landscape

In this section, we explore the constant immunomodulatory parameter $M$, which posits a fixed level of immunomodulatory activity that remains invariant regardless of tumor cell population size. This approach serves as a foundational framework to analyze the interactions between the immune system and tumor cells under static immunomodulatory conditions.

Fig. 6A-B in the main text illustrates the fitted CDF for the constant $M$ immunomodulation, demonstrating its alignment with the clinical data. Additionally, we analyzed the MSE histograms, presented in Figure S13A-B for breast and bladder cancer, respectively. These histograms indicate that the constant $M$ immunomodulation exhibits a strong fit to the clinical data in the early progression stages, as evidenced by the low MSE values. However, as cancer progression advances, the static nature of the constant $M$ model shows limitations, particularly in bladder cancer, where the model struggles to capture the complexity of the disease's progression dynamics. This suggests that while the constant immunomodulation model provides valuable insights, it may not fully account for the adaptive nature of tumor-immune interactions over time.

##### S9.3.2 Dynamic Immunomodulatory Landscape

This section focuses on the dynamic immunomodulatory landscape, specifically addressing the reversible and accumulating immunomodulatory parameters $M$. Unlike the constant $M$ approach, the reversible $M$ case allows for variations in immunomodulatory activity in response to changes in the tumor cell population, effectively capturing the dynamic nature of immune responses and their regulatory effects on tumor growth. The accumulating $M$ case, on the other hand, accounts for a progressively increasing immunomodulatory effect as the tumor evolves, reflecting an intensifying immune suppression over time.

Fig. 6C-D in the main text present the fit of the reversible immunomodulation to clinical data for bladder and breast cancer, respectively, demonstrating its effectiveness in aligning with observed data. These panels also illustrate the temporal changes in $M$, providing a visual representation of the dynamic immunomodulatory activity in these cancer types. Similarly, Fig. 6E-F depict the accumulating immunomodulation for bladder and breast cancer, respectively.

Our analysis of the MSE values, as shown in Fig. S13, reveals that the reversible immunomodulation consistently exhibits a lower average MSE compared to both the constant and accumulating cases. This finding suggests that the reversible immunomodulation provides a superior fit, particularly in the context of breast cancer, where the dynamic adjustments of the immune response more accurately reflect the observed clinical data. The accumulating immunomodulation, while capturing some aspects of the tumor's evolving immune environment, also shows a gradual increase in immunomodulatory strength over time. However, in bladder cancer, all immunomodualtion landscapes exhibit challenges in accurately predicting late-stage progression, highlighting the complexity of immune escape mechanisms in this cancer type. These results underscore the importance of incorporating dynamic elements into models of tumor-immune interactions to better capture the nuances of cancer progression and immune suppression.

### References


[1] Geoffrey Hadfield. The dormant cancer cell. *British medical journal*, 2(4888):607, 1954.

[2] Julio A Aguirre-Ghiso. Models, mechanisms and clinical evidence for cancer dormancy. *Nature Reviews Cancer*, 7(11):834–846, 2007.

[3] Gavin P Dunn, Lloyd J Old, and Robert D Schreiber. The three es of cancer immunoediting. *Annu. Rev. Immunol.*, 22:329–360, 2004.

[4] Deepak Mittal, Matthew M Gubin, Robert D Schreiber, and Mark J Smyth. New insights into cancer immunoediting and its three component phases—elimination, equilibrium and escape. *Current opinion in immunology*, 27:16–25, 2014.

[5] Hao-fan Wang, Sha-sha Wang, Mei-chang Huang, Xin-hua Liang, Ya-Jie Tang, and Ya-ling Tang. Targeting immune-mediated dormancy: a promising treatment of cancer. *Frontiers in oncology*, 9:498, 2019.

[6] Rona M MacKie, Robin Reid, and Brian Junor. Fatal melanoma transferred in a donated kidney 16 years after melanoma surgery. *New England Journal of Medicine*, 348(6):567–568, 2003.

[7] Filippo G Giancotti. Mechanisms governing metastatic dormancy and reactivation. *Cell*, 155(4):750–764, 2013.

[8] Catherine M Koebel, William Vermi, Jeremy B Swann, Nadeen Zerafa, Scott J Rodig, Lloyd J Old, Mark J Smyth, and Robert D Schreiber. Adaptive immunity maintains occult cancer in an equilibrium state. *Nature*, 450(7171):903–907, 2007.

[9] Irene Romero, Cristina Garrido, Ignacio Algarra, Antonia Collado, Federico Garrido, and Angel M Garcia-Lora. T lymphocytes restrain spontaneous metastases in permanent dormancy. *Cancer research*, 74(7):1958–1968, 2014.

[10] Federico Garrido, Natalia Aptsiauri, Elien M Doorduijn, Angel M Garcia Lora, and Thorbald van Hall. The urgent need to recover mhc class i in cancers for effective immunotherapy. *Current opinion in immunology*, 39:44–51, 2016.

[11] Albert C Yeh and Sridhar Ramaswamy. Mechanisms of cancer cell dormancy—another hallmark of cancer? *Cancer research*, 75(23):5014–5022, 2015.

[12] Jean Albrengues, Mario A Shields, David Ng, Chun Gwon Park, Alexandra Ambrico, Morgan E Poindexter, Priya Upadhyay, Dale L Uyeminami, Arnaud Pommier, Victoria Küttner, et al. Neutrophil extracellular traps produced during inflammation awaken dormant cancer cells in mice. *Science*, 361(6409):eaao4227, 2018.

[13] Matthew A Summers, Michelle M McDonald, and Peter I Croucher. Cancer cell dormancy in metastasis. *Cold Spring Harbor Perspectives in Medicine*, 10(4):a037556, 2020.

[14] Kathleen P Wilkie and Philip Hahnfeldt. Mathematical models of immune-induced cancer dormancy and the emergence of immune evasion. *Interface Focus*, 3(4):20130010, 2013.

[15] Kathleen Wilkie, Philip Hahnfeldt, and Lynn Hlatky. Using ordinary differential equations to explore cancer-immune dynamics and tumor dormancy. *bioRxiv*, page 049874, 2016.

[16] Aikaterini Hatziioannou, Themis Alissafi, and Panayotis Verginis. Myeloid-derived suppressor cells and t regulatory cells in tumors: unraveling the dark side of the force. *Journal of leukocyte biology*, 102(2):407–421, 2017.

[17] Amedeo Amedei, Elena Niccolai, Marisa Benagiano, Chiara Della Bella, Fabio Cianchi, Paolo Bechi, Antonio Taddei, Lapo Bencini, Marco Farsi, Paola Cappello, et al. Ex vivo analysis of pancreatic cancer-infiltrating t lymphocytes reveals that eno-specific tregs accumulate in tumor tissue and inhibit th1/th17 effector cell functions. *Cancer Immunology, Immunotherapy*, 62(7):1249–1260, 2013.

[18] Xuefang Cao, Sheng F Cai, Todd A Fehniger, Jiling Song, Lynne I Collins, David R Piwnica-Worms, and Timothy J Ley. Granzyme b and perforin are important for regulatory t cell-mediated suppression of tumor clearance. *Immunity*, 27(4):635–646, 2007.

[19] Theresa L Whiteside. The role of regulatory t cells in cancer immunology. *ImmunoTargets and therapy*, pages 159–171, 2015.

[20] Ralf-Peter Czekay, Dong-Joo Cheon, Rohan Samarakoon, Stacie M Kutz, and Paul J Higgins. Cancer-associated fibroblasts: mechanisms of tumor progression and novel therapeutic targets. *Cancers*, 14(5):1231, 2022.

[21] Raisa A Glabman, Peter L Choyke, and Noriko Sato. Cancer-associated fibroblasts: Tumorigenicity and targeting for cancer therapy. *Cancers*, 14(16):3906, 2022.

[22] Giovanna Angelini, Stefania Gardella, Massimo Ardy, Maria Rosa Ciriolo, Giuseppe Filomeni, Giovanna Di Trapani, Frank Clarke, Roberto Sitia, and Anna Rubartelli. Antigen-presenting dendritic cells provide the reducing extracellular microenvironment required for t lymphocyte activation. *Proceedings of the National Academy of Sciences*, 99(3):1491–1496, 2002.

[23] Minu K Srivastava, Pratima Sinha, Virginia K Clements, Paulo Rodriguez, and Suzanne Ostrand-Rosenberg. Myeloid-derived suppressor cells inhibit t-cell activation by depleting cystine and cysteine. *Cancer research*, 70(1):68–77, 2010.

[24] Ngozi R Monu and Alan B Frey. Myeloid-derived suppressor cells and anti-tumor t cells: a complex relationship. *Immunological investigations*, 41(6-7):595–613, 2012.

[25] Bo Huang, Ping-Ying Pan, Qingsheng Li, Alice I Sato, David E Levy, Jonathan Bromberg, Celia M Divino, and Shu-Hsia Chen. Gr-1+ cd115+ immature myeloid suppressor cells mediate the development of tumor-induced t regulatory cells and t-cell anergy in tumor-bearing host. *Cancer research*, 66(2):1123–1131, 2006.

[26] Cunren Liu, Shaohua Yu, John Kappes, Jianhua Wang, William E Grizzle, Kurt R Zinn, and Huang-Ge Zhang. Expansion of spleen myeloid suppressor cells represses nk cell cytotoxicity in tumor-bearing host. *Blood*, 109(10):4336–4342, 2007.

[27] Jason T George and Herbert Levine. Stochastic modeling of tumor progression and immune evasion. *Journal of theoretical biology*, 458:148–155, 2018.

[28] Jason T George and Herbert Levine. Sustained coevolution in a stochastic model of cancer–immune interaction. *Cancer research*, 80(4):811–819, 2020.

[29] Jason T George and Herbert Levine. Implications of tumor–immune coevolution on cancer evasion and optimized immunotherapy. *Trends in Cancer*, 7(4):P373–383, 2021.

[30] Jason T George, David A Kessler, and Herbert Levine. Effects of thymic selection on t cell recognition of foreign and tumor antigenic peptides. *Proceedings of the National Academy of Sciences*, pages E7875–E7881, 2017.

[31] Samuel Karlin. *A first course in stochastic processes.* Academic press, 2014.

[32] Stewart N Ethier and Thomas G Kurtz. *Markov processes: characterization and convergence.* John Wiley & Sons, 2009.

[33] Thomas G Kurtz and Philip Protter. Weak limit theorems for stochastic integrals and stochastic differential equations. *The Annals of Probability*, pages 1035–1070, 1991.

[34] Wai-Yuan Tan. *Stochastic models with applications to genetics, cancers, AIDS and other biomedical systems*, volume 19. World Scientific, 2015.

[35] Nicole E Scharping, Ashley V Menk, Rebecca S Moreci, Ryan D Whetstone, Rebekah E Dadey, Simon C Watkins, Robert L Ferris, and Greg M Delgoffe. The tumor microenvironment represses t cell mitochondrial biogenesis to drive intratumoral t cell metabolic insufficiency and dysfunction. *Immunity*, 45(2):374–388, 2016.

[36] McLane J Watson, Paolo DA Vignali, Steven J Mullett, Abigail E Overacre-Delgoffe, Ronal M Peralta, Stephanie Grebinoski, Ashley V Menk, Natalie L Rittenhouse, Kristin DePeaux, Ryan D Whetstone, et al. Metabolic support of tumour-infiltrating regulatory t cells by lactic acid. *Nature*, 591(7851):645–651, 2021.

[37] Kuhn Lab at USC. Breast cancer progression models. `https://kuhn.usc.edu/breast_cancer/`.

[38] Paul K. Newton, Jeremy Mason, Neethi Venkatappa, Maxine S. Jochelson, Brian Hurt, Jorge Nieva, Elizabeth Comen, Larry Norton, and Peter Kuhn. Spatiotemporal progression of metastatic breast cancer: a markov chain model highlighting the role of early metastatic sites. *npj Breast Cancer*, 1(1):15018, Oct 2015.

[39] Kuhn Lab at USC. Bladder cancer progression models. `https://kuhn.usc.edu/bladder_cancer/`.

[40] Zaki Hasnain, Jeremy Mason, Karanvir Gill, Gus Miranda, Inderbir S. Gill, Peter Kuhn, and Paul K. Newton. Machine learning models for predicting post-cystectomy recurrence and survival in bladder cancer patients. *PLOS ONE*, 14(2):1–15, 02 2019.
